# Bacterial DNAemia is associated with serum zonulin levels in older subjects

**DOI:** 10.1101/2020.04.10.035519

**Authors:** Giorgio Gargari, Valentina Taverniti, Cristian Del Bo’, Stefano Bernardi, Cristina Andres-Lacueva, Raul González-Domínguez, Paul A. Kroon, Mark S. Winterbone, Antonio Cherubini, Patrizia Riso, Simone Guglielmetti

**Author notes:** Corresponding author. Mailing address: Via Celoria 2, 20133, Milano, Italia.

## Abstract

The increased presence of bacteria in blood is a plausible contributing factor in the development and progression of aging-associated diseases. In this context, we performed the quantification and the taxonomic profiling of the bacterial DNA in blood samples collected from a group of forty-three older subjects enrolled in a nursing home. Quantitative PCR targeting the 16S rRNA gene revealed that all the older volunteers contained detectable amounts of bacterial DNA in their blood. The total amount of 16S rRNA gene copies varied considerably between subjects. Correlation analyses revealed that the bacterial DNAemia (expressed as concentration of 16S rRNA gene copies in blood) significantly correlated with the serum levels of zonulin, an emerging marker of intestinal permeability. This result was confirmed by the analysis of a second set of blood samples collected after approximately four months from the same subjects. Analyses of 16S rRNA gene profiling revealed that most of the bacterial DNA detected in blood was ascribable to the phylum *Proteobacteria* with a predominance of *Pseudomonadaceae* and *Enterobacteriaceae*. Several control samples were also analyzed to assess the influence exerted by contaminant bacterial DNA potentially originating from reagents and materials. The date reported here suggest that para-cellular permeability of epithelial (and potentially also endothelial) cell layers may play an important role in bacterial migration into the bloodstream. Bacterial DNAemia is likely to impact on several aspects of host physiology and could underpin the development and prognosis of various diseases in older subjects.

## Introduction

The advent and diffusion of metagenomic approaches has confuted the dogma that several tissues of the human body were completely aseptic, such as amniotic fluid (Collado et al 2016), the bladder (Thomas-White et al 2016), and blood (Paisse et al 2016). In particular, several studies in the last decade reported the existence of bacterial DNA in the bloodstream of healthy individuals (Paisse et al 2016). For instance, in 2001, Nikkari et al. reported the presence of significant amounts of bacterial DNA in venous blood specimens from four healthy subjects, and hypothesized the existence of a “normal” population of hematic bacterial DNA molecules (Nikkari et al 2001). Nonetheless, the precise consequences on host physiology of such bacterial DNAemia are still largely undefined and potentially underestimated (Gosiewski et al 2017, Potgieter et al 2015).

Bacterial DNA in blood plausibly originates from (i) direct translocation of microbial cells through an injured epithelial barrier, (ii) microbe sampling activity of antigen-presenting cells (dendritic cells, macrophages) at the epithelium, or (iii) via microfold (M) cells in Peyer’s patches of the intestine (Potgieter et al 2015). These events increase the presence of bacterial factors found systemically (in bloodstreams and organs), resulting in a significant stimulation of the host’s immune system. In line with this assumption, blood microbiota abundance and composition were reported to be associated with chronic, inflammatory diseases (Potgieter et al 2015) and non-communicable diseases such as type II diabetes (Amar et al 2011, Sato et al 2014) and cardiovascular disease (Amar et al 2013). Indeed, the presence / size of a blood microbiota population was proposed as a biomarker for the assessment of cardiovascular risk (Amar et al 2013). Overall, the experimental evidence suggests that higher levels of bacterial DNA in blood may be indicative of increased risk of disease, particularly for subjects characterized by other concomitant risk factors, such as older subjects.

Aging is typically characterized by chronic low-grade systemic inflammation (inflammaging; (Franceschi et al 2000)), and it’s severity was reported to be a valid predictor of overall illness and mortality in an older population (Franceschi and Campisi 2014). Thevaranjan et al. demonstrated in a mouse model that age-related microbial dysbiosis increases intestinal permeability, permitting the translocation into the bloodstream of microbial products that, consequently, trigger systemic inflammation (Thevaranjan et al 2017)). Therefore, this report provides evidence that supports the notion of a mechanistic link between bacterial DNAemia and inflammaging. In addition, it supports the hypothesis that intestinal barrier deterioration occurs during aging, promoting increased absorption of luminal factors (Ma et al 1992), and supports the now more than 100-year-old Metchnikoff speculation that, during aging, “*the intestinal microbes or their poisons may reach the system generally and bring harm to it*” (Metchnikoff 1907).

Starting from the notion that bacterial atopobiosis (i.e., “*microbes that appear in places other than their normal location*”, (Potgieter et al 2015)) in blood has a significant role in the development and progression of aging and aging-associated diseases, we performed the quantification and the taxonomic profiling of the bacterial DNA in blood collected from a group of forty-three older subjects. The data have been correlated with several metabolic and functional markers, including zonulin as marker of intestinal permeability. To the best of our knowledge, this is the first report of the amounts and sources of bacterial DNA in the peripheral blood of a group of older subjects.

## Subjects and Methods

### Older population

This study involved the analysis of samples obtained from older volunteers (n=43; 28 women and 15 men; average age: 79.2 ± 9.8 y; body mass index (BMI): 26.5 ± 5.8; **Table 1**) enrolled within the project entitled “*Gut and blood microbiomics for studying the effect of a polyphenol-rich dietary pattern on intestinal permeability in the elderly*” (MaPLE project). Subjects were recruited in a well-controlled setting at Civitas Vitae (OIC Foundation, Padua, Italy which include residential care and independent residences for older subjects) based on specific inclusion and exclusion criteria as previously reported in detail (Guglielmetti et al 2020).

**Table 1.**
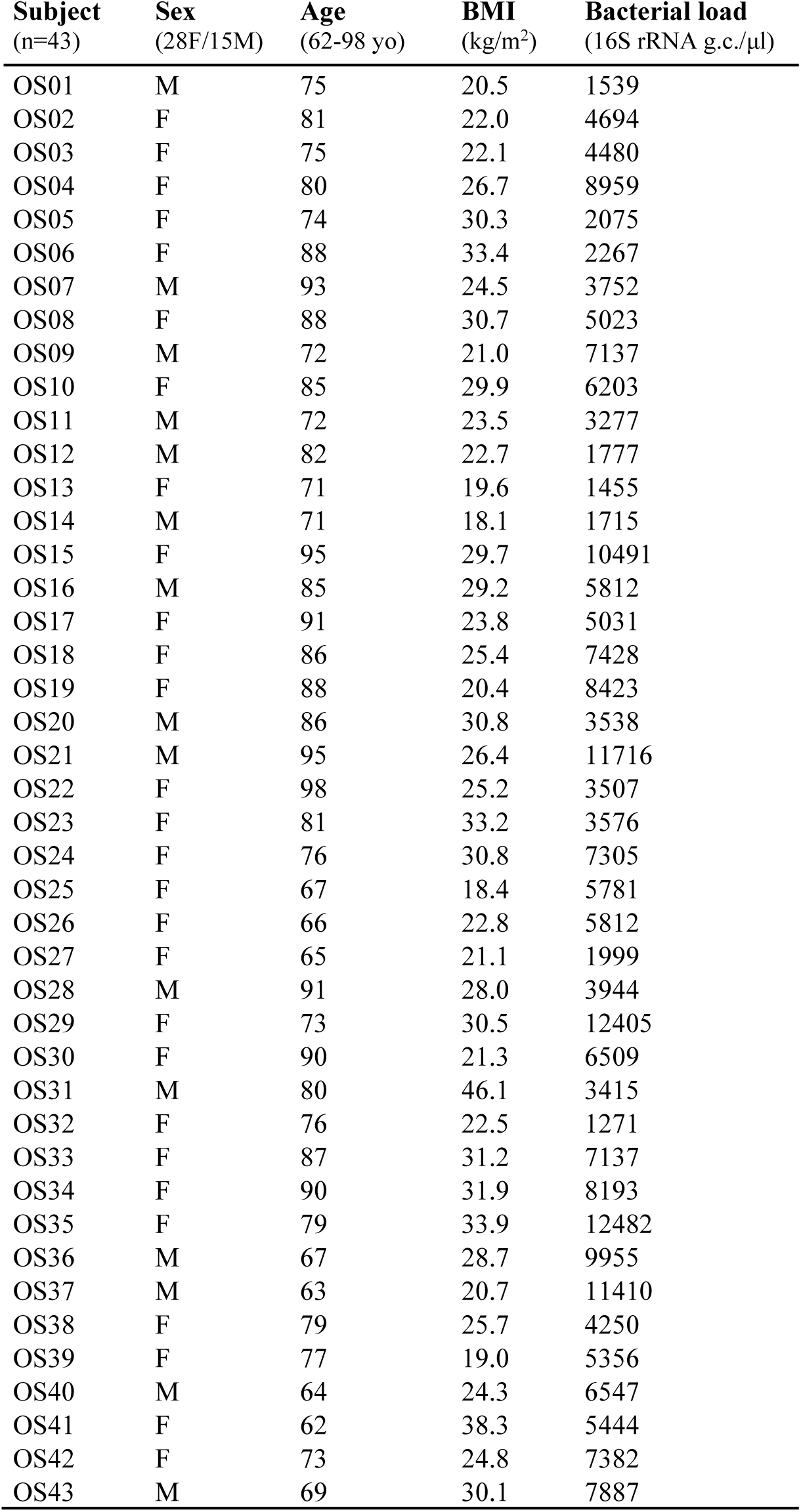
Demographic and anthropometrical characteristics of older volunteers recruited for the study. Blood bacterial load (determined by qPCR and expressed as 16S rRNA gene copies per μl) is also shown. OS01-43, consecutive anonymized coding of the older subjects included in the study.16S rRNA g.c., 16S rRNA gene copies.

| **Subject**<br>(n=43) | **Sex**<br>(28F/15M) | **Age**<br>(62-98 yo) | **BMI**<br>($\text{kg}/\text{m}^2$) | **Bacterial load**<br>(16S rRNA g.c./$\mu\text{l}$) |
| --- | --- | --- | --- | --- |
| OS01 | M | 75 | 20.5 | 1539 |
| OS02 | F | 81 | 22.0 | 4694 |
| OS03 | F | 75 | 22.1 | 4480 |
| OS04 | F | 80 | 26.7 | 8959 |
| OS05 | F | 74 | 30.3 | 2075 |
| OS06 | F | 88 | 33.4 | 2267 |
| OS07 | M | 93 | 24.5 | 3752 |
| OS08 | F | 88 | 30.7 | 5023 |
| OS09 | M | 72 | 21.0 | 7137 |
| OS10 | F | 85 | 29.9 | 6203 |
| OS11 | M | 72 | 23.5 | 3277 |
| OS12 | M | 82 | 22.7 | 1777 |
| OS13 | F | 71 | 19.6 | 1455 |
| OS14 | M | 71 | 18.1 | 1715 |
| OS15 | F | 95 | 29.7 | 10491 |
| OS16 | M | 85 | 29.2 | 5812 |
| OS17 | F | 91 | 23.8 | 5031 |
| OS18 | F | 86 | 25.4 | 7428 |
| OS19 | F | 88 | 20.4 | 8423 |
| OS20 | M | 86 | 30.8 | 3538 |
| OS21 | M | 95 | 26.4 | 11716 |
| OS22 | F | 98 | 25.2 | 3507 |
| OS23 | F | 81 | 33.2 | 3576 |
| OS24 | F | 76 | 30.8 | 7305 |
| OS25 | F | 67 | 18.4 | 5781 |
| OS26 | F | 66 | 22.8 | 5812 |
| OS27 | F | 65 | 21.1 | 1999 |
| OS28 | M | 91 | 28.0 | 3944 |
| OS29 | F | 73 | 30.5 | 12405 |
| OS30 | F | 90 | 21.3 | 6509 |
| OS31 | M | 80 | 46.1 | 3415 |
| OS32 | F | 76 | 22.5 | 1271 |
| OS33 | F | 87 | 31.2 | 7137 |
| OS34 | F | 90 | 31.9 | 8193 |
| OS35 | F | 79 | 33.9 | 12482 |
| OS36 | M | 67 | 28.7 | 9955 |
| OS37 | M | 63 | 20.7 | 11410 |
| OS38 | F | 79 | 25.7 | 4250 |
| OS39 | F | 77 | 19.0 | 5356 |
| OS40 | M | 64 | 24.3 | 6547 |
| OS41 | F | 62 | 38.3 | 5444 |
| OS42 | F | 73 | 24.8 | 7382 |
| OS43 | M | 69 | 30.1 | 7887 |

### Blood sampling

Blood drawing from a peripheral vein was performed early in the morning from fasted subjects. Samples were immediately processed; 0.5 ml aliquots of whole blood were stored at -80°C until analysis of blood bacterial DNA in certified DNase/RNase free and pyrogen free vials, while the remaining blood was centrifuged at 1000 x g for 15 min at room temperature and the obtained serum stored at -80°C for the subsequent analysis of metabolic/functional markers.

### DNA extraction, qPCR experiments and sequencing of 16S rRNA gene amplicons

Bacterial DNA quantification and sequencing reactions were performed by Vaiomer SAS (Labège, France) using optimized blood-specific techniques as described earlier (Lelouvier et al 2016, Paisse et al 2016). Since low microbial biomass was expected, four technical controls (ultrapure water (UPW) samples) were analyzed in the same way as the blood samples (C_extr_). Two additional control samples (C_EDTA_) were prepared by adding 2 ml of commercial phosphate buffered saline (PBS; Sigma-Aldrich) into an EDTA tube, directly (C_dEDTA_) or after passage through a vacutainer blood collection tube (C_vEDTA_). In brief, DNA was extracted from 100 µl of whole blood or control samples and quantified by quantitative real-time PCR (qPCR) experiments using bacterial primers EUBF 5’-TCCTACGGGAGGCAGCAGT-3’ and EUBR 5’-GGACTACCAGGGTATCTAATCCTGTT-3’ (Nadkarni et al 2002), which target the V3-V4 hypervariable regions of the bacterial 16S rRNA gene with 100% specificity (i.e., no eukaryotic, mitochondrial, or Archaea DNA is targeted) and high sensitivity (16S rRNA of more than 95% of bacteria in Ribosomal Database Project database are amplified). Four samples of UPW were used as templates in qPCR reactions for technical control (C_qPCR_). The 16S rRNA gene was quantified by qPCR in triplicate and normalized using a plasmid-based standard scale. Finally, the results were reported as number of copies of 16S rRNA gene per µl of blood.

DNA extracted from whole blood and the four C_qPCR_ controls were also used for 16S rRNA gene taxonomic profiling using MiSeq Illumina technology (2 × 300 paired-end MiSeq kit V3, set to encompass 467-bp amplicon) according to specific protocol and primers from Vaiomer as previously described (Lluch et al 2015, Paisse et al 2016). Then, sequences were analyzed using Vaiomer bioinformatic pipeline to determine bacterial community profiles. Briefly, after demultiplexing of the bar-coded Illumina paired reads, single read sequences were trimmed (reads R1 and R2 to respectively 290 and 240 bases) and paired for each sample independently into longer fragments, non-specific amplicons were removed and remaining sequences were clustered into operational taxonomic units (OTUs, here also called “Cluster”) using FROGS v1.4.0 (Escudie et al 2018) with default parameters. A taxonomic assignment was performed against the Silva 128 Parc database, and diversity analyses and graphical representations were generated using the R PhyloSeq v1.14.0 package. In addition, the 16S rRNA gene profiling analysis of C_dEDTA_ and C_vEDTA_ control samples was carried out in an independent experiment together with additional four extraction controls (C_extr_2) and an additional blood sample collected from one of the older volunteers. For bacterial taxonomic profiling and correlation analyses, the relative abundance of each taxon was normalized against the total number 16S rRNA gene copies in that specific sample.

### Blood parameters analyses

Fasting serum levels of glucose, insulin, creatinine, uric acid, aspartate aminotransferase (AST), alanine aminotransferase (ALT), and gamma-glutamyl transferase (GGT), triglyceride (TG), total cholesterol (TC), low-density lipoprotein cholesterol (LDL-C) were determined by an automatic biochemical analyzer (ILAB650, Instrumentation Laboratory and TOSO AIA 900, Italy), while the high-density lipoprotein cholesterol (HDL-C) concentration was estimated using the Friedewald formula (Guglielmetti et al 2020). The homeostatic model assessment (HOMA) index was calculated with the formula HOMA-IR= [fasting glucose (mg/dL) × insulin (IU)/405]. Serum zonulin levels were quantified using IDK® Zonulin ELISA Kit (Immundiagnostik AG, Germany). C-reactive protein (CRP) was quantified using the Quantikine Human C-reactive protein ELISA kit (R&D Systems cat# DCRP00), TNFα using the Quantikine high sensitivity Human TNF-α ELISA kit (R&D Systems cat# HSTA00E) and IL6 using the Quantikine high sensitivity Human IL-6 ELISA kit (R&D Systems cat# HS600B). Intercellular adhesion molecule-1 (ICAM-1) and vascular cell adhesion molecule-1 (VCAM-1) were quantified in serum samples with an ELISA kit (Booster^®^ from Vinci Biochem S.r.l., Vinci, Italy). The Cockcroft-Gault equation was used for estimating creatinine clearance (CG index).

### Statistics

Statistical analyses were performed using R statistical software (version 3.1.2). The correlation analyses were carried out using the Kendall and Spearman analysis, as previously described (Gargari et al 2018a, Gargari et al 2018b), calculated between blood bacterial load, age, BMI, and clinical and molecular parameters quantified in the blood of older volunteers at recruitment. The regression analysis was performed to evaluate the linear relationship between total bacteria load and zonulin levels, in addition to Pearson’s correlation analysis. Statistical significance was set at P ≤ 0.05; differences with 0.05 < P ≤ 0.10 were accepted as trends. When P values correction was applied, the Holm-Bonferroni method was used.

## Results

### Older subjects harbor bacterial DNA in blood

qPCR assays with pan-bacterial primers targeting the 16S rRNA gene were used to detect and quantify DNA of bacterial origin in the blood samples collected from 43 older people. In addition, the same analysis was performed on 10 control samples: Extraction controls (C_extr_; four samples), EDTA tube and vacutainer system controls (C_dEDTA_ and C_vEDTA_; two samples) and qPCR controls (C_qPCR_ controls; four samples). Bacterial DNA was detected in all blood samples collected from the older subjects under study. Specifically, we quantified a minimum of 1271 and a maximum of 12482 copies of the 16S rRNA gene per µl of blood (Table 1 and **Fig. 1**). Conversely, the gene copies detected in controls were much lower than those found in blood samples, with a mean detection (± standard deviation) of 156 (± 29), 69 (± 3) and 252 (± 190) 16S rRNA gene copies per equivalent μl of blood for C_extr_, C_EDTA_, and C_qPCR_ samples, respectively (**Fig. 1**).

**Fig. 1.**
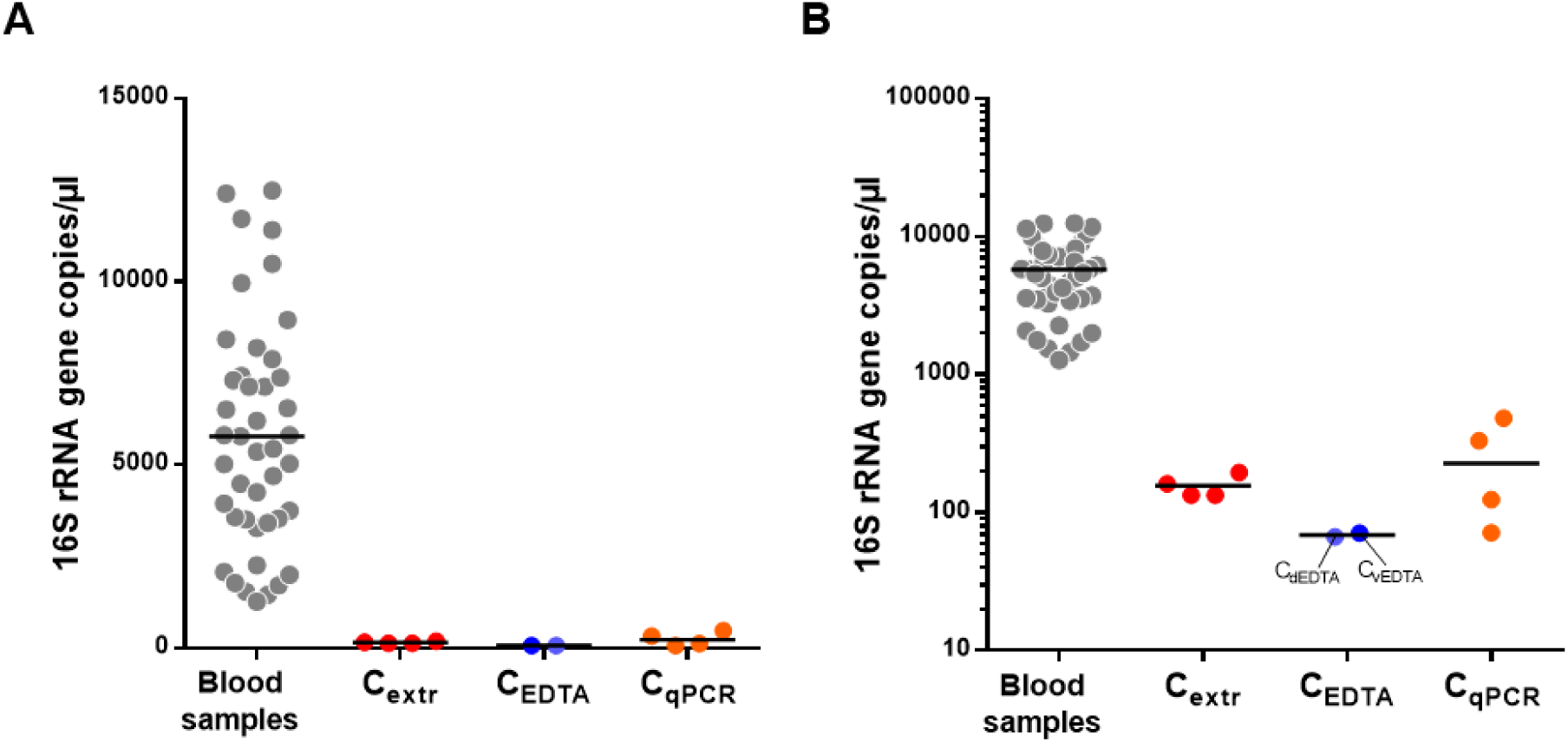
Bacterial load in blood samples of older volunteers (n=43) and in technical controls as determined by qPCR with panbacterial primers targeting the 16S rRNA gene. Data are reported as copies of the gene per volume of blood in linear (**A**) and logarithmic (**B**) scale. C_extr_, DNA extracted from ultrapure deionized water with the same protocol used for blood samples; C_EDTA_, DNA extracted with the same protocol used for blood samples from phosphate buffered saline (PBS) into an EDTA tube, added directly (C_dEDTA_) or after passage through a vacutainer blood collection tube (C_vEDTA_); C_qPCR_, ultrapure deionized water directly used as template in qPCR reactions. Horizontal black lines indicate the mean.

Overall, these results indicate that the older volunteers under study harbored detectable amounts of bacterial DNA in their blood, in a concentration that varied approximately of one order of magnitude among samples.

### The amount of bacterial DNA in blood of older subjects is associated with serum zonulin

In the subsequent part of the study, we performed correlation analyses between the blood bacterial DNA load and several clinical and molecular parameters that are relevant for metabolic and physiological function in the older population. Specifically, besides age and BMI, markers related to immune status (serum CRP, IL6, and TNFα concentrations), vascular function (serum ICAM-1 and VCAM-1), glucose metabolism/homeostasis (serum glucose, insulin, and the derived HOMA index), renal function (urinary creatinine, uric acid, and CG index), lipid metabolism (serum TC, HDL-C, LDL-C, and TG), liver function (serum AST, ALT, and GGT) and epithelial permeability (serum zonulin) were also quantified and possible associations with blood bacterial DNA load investigated. A number of expected strong associations were observed, for example between TC and LDL cholesterol, insulin and HOMA index, IL6 and CRP, and GOT (AST) and GPT (ALT) (**Fig. 2**). In addition, we found clear inverse correlations of the CG index with creatinine and age, and between GOT (AST) and CRP. We also found a significant correlation between the 16S rRNA gene copy number and HOMA index, and between zonulin and triglycerides. However, in the context of the present study, the most interesting result was the significant positive association between blood bacterial DNA and zonulin (Fig. 2). Specifically, the 16S rRNA gene copy number was found to significantly correlate (Spearman’s ρ = 0.504; Kendall’s P = 0.0004) and linearly associate (Pearson’s r = 0.581; P < 0.0001) with serum zonulin levels (Fig. 2 and **Fig. 3**).

**Fig. 2.**
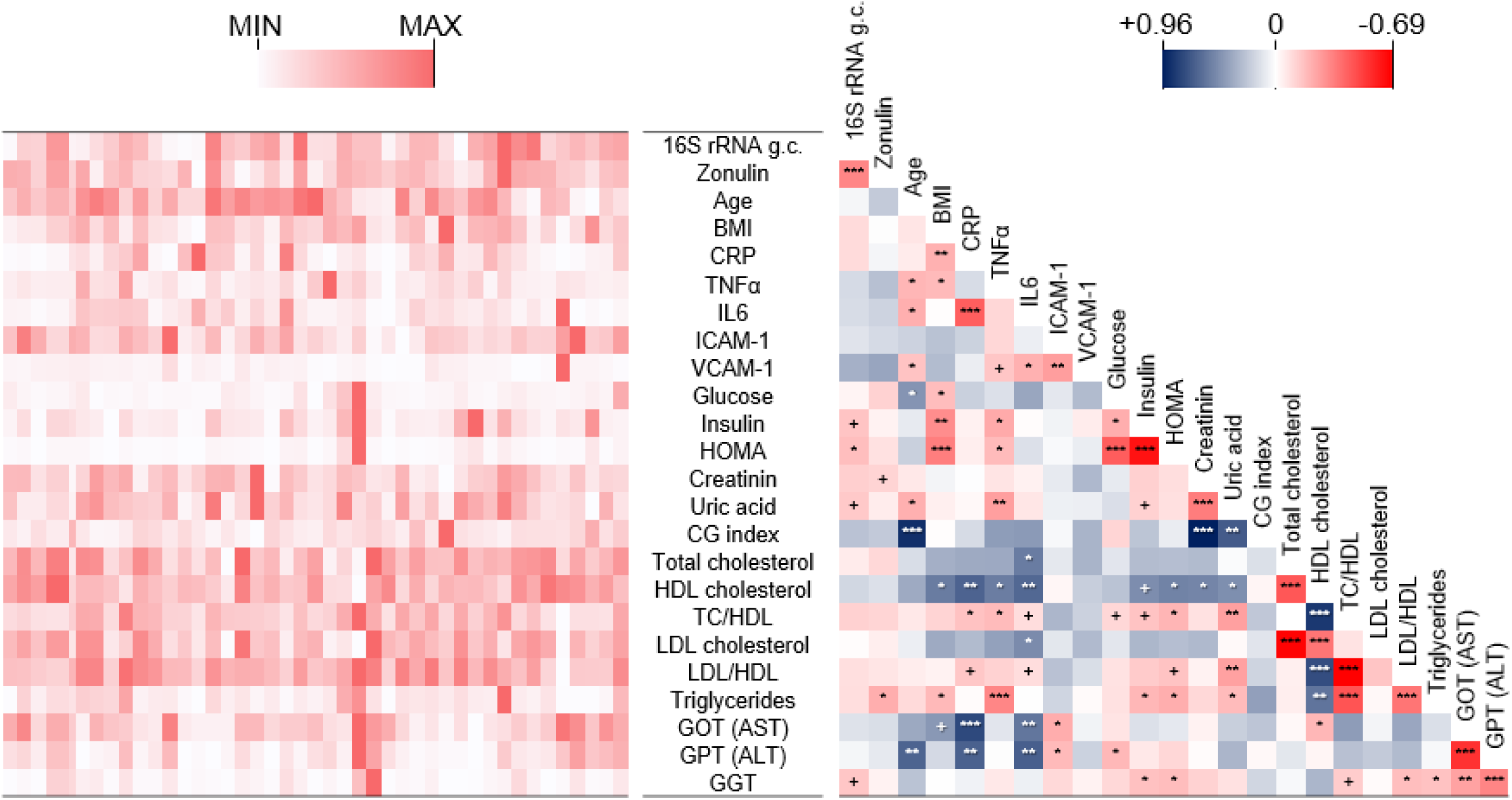
Correlations between bacterial DNAemia, age, BMI, and metabolic and functional markers determined in blood of the older subjects under study (n=43). The heatmaps on the left (white-red color gradient) refers to the distribution among older subject of the item reported in the central column. The heatmap on the right (blue-white-red color gradient) represents the Spearman’s correlation coefficient, ρ. Asterisks indicate the Kendall rank correlation P value: ***, P<0.001; **, P<0.01; *, P<0.05; +, 0.05<P≤0.10.

**Fig. 3.**
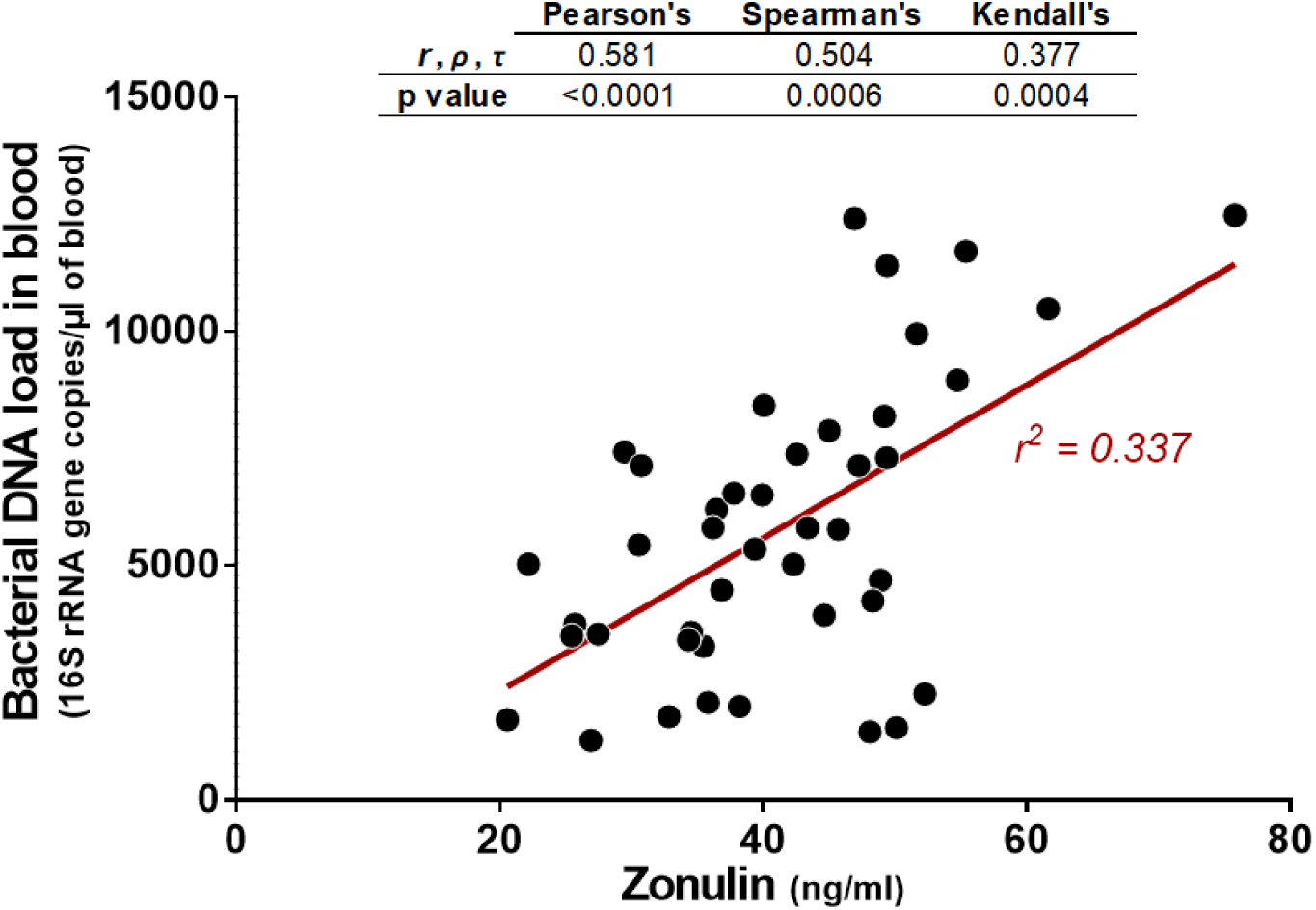
Linear regression and correlation analysis carried out between blood bacterial load and serum zonulin levels. Correlation has been assessed through Pearson’s (r coefficient), Spearman’s (ρ), and Kendall’s (τ) analyses.

In order to confirm the observed direct correlation between bacterial DNA load in peripheral blood and serum zonulin levels, blood was collected again from the same cohort of older subjects after approximately four months. Then, 16S rRNA gene copies and zonulin levels were quantified and the results used for correlation and regression analyses. One sample was removed from the analyses due to technical problems during zonulin quantification (total samples considered, n=42). The obtained results revealed again the significant correlation (Spearman’s ρ = 0.486; Kendall’s P = 0.001) and linear association (Pearson’s r = 0.407; P = 0.008) between 16S rRNA gene copies and zonulin levels in blood samples (**Fig. S1**).

Finally, the comparison of results obtained from blood samples collected before and after 4 months revealed a higher inter-subject than intra-subject variability for zonulin and bacterial DNA abundances. In fact, we found a significant correlation and linear association between the two set of samples for both bacterial DNA load (Spearman’s ρ = 0.379; Kendall’s P = 0.013; Pearson’s P = 0.010) zonulin levels (Spearman’s ρ = 0.526; Kendall’s P < 0.001; Pearson’s P = 0.001).

Overall, these results indicate that circulating bacterial DNA is stably linked with serum zonulin level in the group of relatively healthy older people that were studied.

### Taxonomic profiling of blood bacterial DNA

The same blood DNA samples used in qPCR experiments (n=43) were also used in 16S rRNA gene profiling analysis to define their bacterial taxonomic composition. In addition, in the same analysis, we also taxonomically profiled four C_qPCR_ samples, since they showed the highest concentration of 16S rRNA gene copies among controls (Fig. 1). The average number of reads assigned to OTUs was around 35,000 per sample. Curve plots for rarefaction analysis suggested that the sequence depth captured the diversity in all samples (**Fig. S2A**). Notably, the OTU richness of blood samples (mean ± standard deviation = 40 ± 8 OTUs per sample) was significantly higher (twice, on average) than the richness of the controls (mean ± standard deviation = 21 ± 2 OTUs per sample) (**Fig. S2B**). In addition, multidimensional scaling (MDS) representation of β-diversity calculated through generalized UniFrac distance clustered controls outside the group of blood samples (**Fig. S2C**).

Subsequently, we analyzed the bacterial composition of blood DNA samples at different taxonomic levels. In order to infer information concerning taxonomic abundance, we normalized the number of reads by multiplying the relative abundance of taxa by the 16S rRNA gene abundance determined by qPCR in each blood sample. This analysis revealed that the phylum *Proteobacteria* was the most represented in all samples, followed by *Actinobacteria*. These two phyla were found in all samples, whereas *Firmicutes* and *Bacteroidetes* were found in 98% and 79% of subjects, respectively (**Fig. 4A**). *Fusobacteria* were sporadically detected (in about 28% of samples), whereas other phyla were found in less than 5% of samples. Only three families, i.e. *Enterobacteriaceae, Pseudomonadaceae* (phylum *Proteobacteria*), and *Micrococcaceae* (phylum *Actinobacteria*) were found in all samples (**Fig. 4B**). *Pseudomonas* was the most abundant genus in all samples except one, in which the *Proteobacteria* genus *Massilia* was predominant. The second most abundant genus was *Arthrobacter* (family *Micrococcaceae*), followed by the *Proteobacteria* genera *Phyllobacterium* (family *Phyllobacteriaceae*), *Acinetobacter* (family *Moraxellaceae*), *Paracoccus* (family *Rhodobacteraceae*), and *Tepidimonas* (family *Comamonadaceae*) (**Fig. 4C**).

**Fig. 4.**
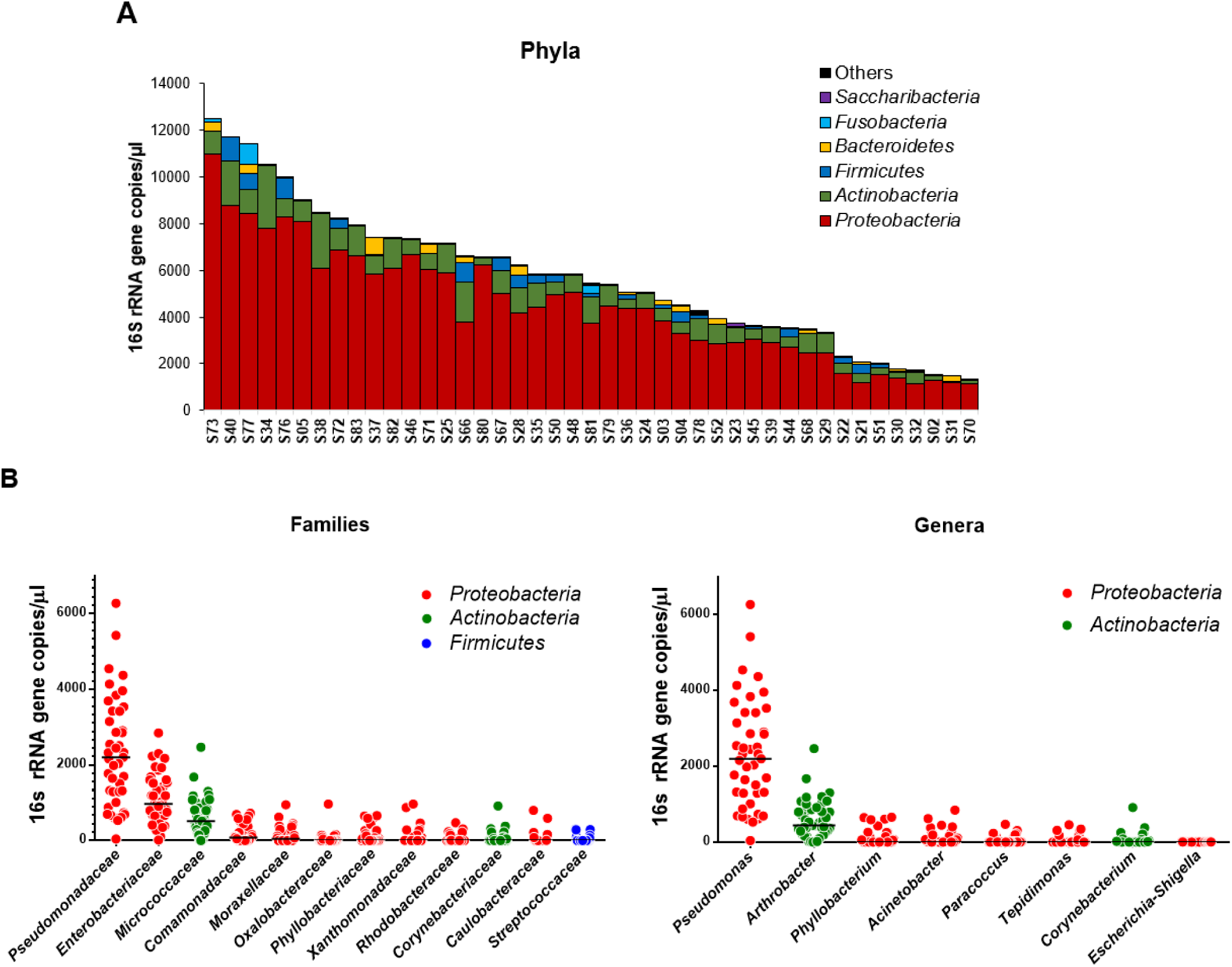
Taxonomic composition of the bacterial DNA detected in the blood of older subjects expressed as 16S rRNA gene copies per volume of blood. **A**, distribution of phyla in each analyzed blood sample. **B**, most abundant families and genera detected in the blood samples under study.

Finally, the most abundant OTUs (i.e., having a median abundance > 100 16S rRNA gene copies per μl of blood; clusters 1 to 4) were further taxonomically investigated by manual searching against the 16S rRNA gene database within the GenBank by Basic Local Alignment Search Tool (BLAST). The DNA sequence characterizing Cluster 1 showed 100% similarity with species *Pseudomonas fluorescens* and *Pseudomonas canadensis*. Cluster 2 showed 100% similarity with several *Enterobacteriaceae* species (e.g. *Brennaria alni, Escherichia coli, Escherichia fergusonii, Pseudescherichia vulneris, Shigella flexneri, Shigella sonnei*). Cluster 3 was 100% similar to *Pseudomonas azotoformans, Pseudomonas lactis*, and *Pseudomonas paralactis*, which are species within the *P. fluorescens* species group (von Neubeck et al 2017). Finally, the highest match for Cluster 4 was found with the species *Arthrobacter russicus* (99.76%).

Subsequently, in order to find potential contaminants, the taxonomic composition of the DNA from C_qPCR_ samples was analyzed and the abundance of bacterial families and genera was compared between blood and controls. After normalization to 16S rRNA gene copies, the abundance of bacterial taxa in control samples was much lower than that of blood samples (**Fig. S3A-B**). Consequently, we compared the bacterial taxa in blood *vs* control samples without normalization, i.e. using the relative abundance (percentages) (**Fig. S3C-D**). This comparison revealed comparable relative abundance between controls and blood samples only for families *Enterobacteriaceae* and *Micrococcaceae* and the genera *Arthrobacter* and *Escherichia*/*Shigella*. In addition, *Moraxellaceae* and the genus *Acinetobacter* were higher compared to blood samples for only one of the four control samples. Finally, also *Pseudomonadaceae* and the genus *Pseudomonas* were detected in controls, but in lower percentages than largely most of the blood samples (**Fig. S3**).

To find potential contaminants deriving from any reagent and material used in the analysis, successively, we also taxonomically profiled by 16S rRNA gene sequencing four C_extr_ control samples, and the two C_EDTA_ controls together with one blood sample collected from an older volunteer. Again, the amount of bacterial DNA detected in control samples was much lower than that of the blood sample (**Fig. S4A**). About 30 thousand sequencing reads were generated per samples, apart from sample C_dEDTA_, for which approximately 13000 were obtained (**Fig. S4B**). Analysis of sequencing reads indicated a wide taxonomic variability between the control samples, whereas the blood samples comprised a bacterial community structure resembling that observed in the older subjects’ blood samples previously analyzed (**Fig. S4C-D**), which was characterized by *Pseudomonas* as the most abundant genus. The second most abundant bacterial group in the blood sample was ascribed to *Escherichia*/*Shigella*, and it was noted that this taxonomic unit was similarly abundant, in percentage terms, in several control samples. All the other taxonomic units found with a relative abundance higher than 1% in the blood sample were not detected in controls or found in very low concentrations (**Fig. S4C-D**).

Overall, these results strongly suggest that the bacterial DNA detected in the peripheral blood of older subjects is mostly ascribable to the Gram-negative phylum *Proteobacteria*, with the predominance of the genus *Pseudomonas*.

### Serum zonulin is associated to specific bacterial taxa in blood

We performed correlation analyses between the same host markers described above and the taxonomic composition of blood bacterial DNA expressed as abundances normalized to 16S rRNA gene copies. Forty-seven taxa significantly correlated with at least one of the parameters considered (**Fig. 5**), about three quarters of which (33 taxa) belonging to the phylum *Proteobacteria*. The most numerous correlations with bacterial taxa were found for zonulin. In specific, seventeen bacterial taxa showed a significant positive correlation with zonulin levels, whereas none was inversely correlated. Zonulin levels mostly correlated with taxa within the class *γ-Proteobacteria*, i.e. the families *Enterobacteriaceae* and *Pseudomonadaceae*, and the genera *Pseudomonas* and *Escherichia/Shigella* (**Fig. 5**). In addition, a positive correlation with zonulin was observed for the phylum *Actinobacteria* and the class *Actinobacteria*, and for the class *γ-Proteobacteria* and the order Rhizobiales. The correlation between zonulin levels and the abundance of the phyla *Actinobacteria* and *Proteobacteria*, the order *γ-Proteobacteria*, the family *Pseudomonadaceae* and the genus *Pseudomonas* was also confirmed by the analysis of data generated by 16S rRNA gene profiling of the second set of blood samples collected from the older volunteers (n=42) (**Fig. S5**).

**Fig. 5.**
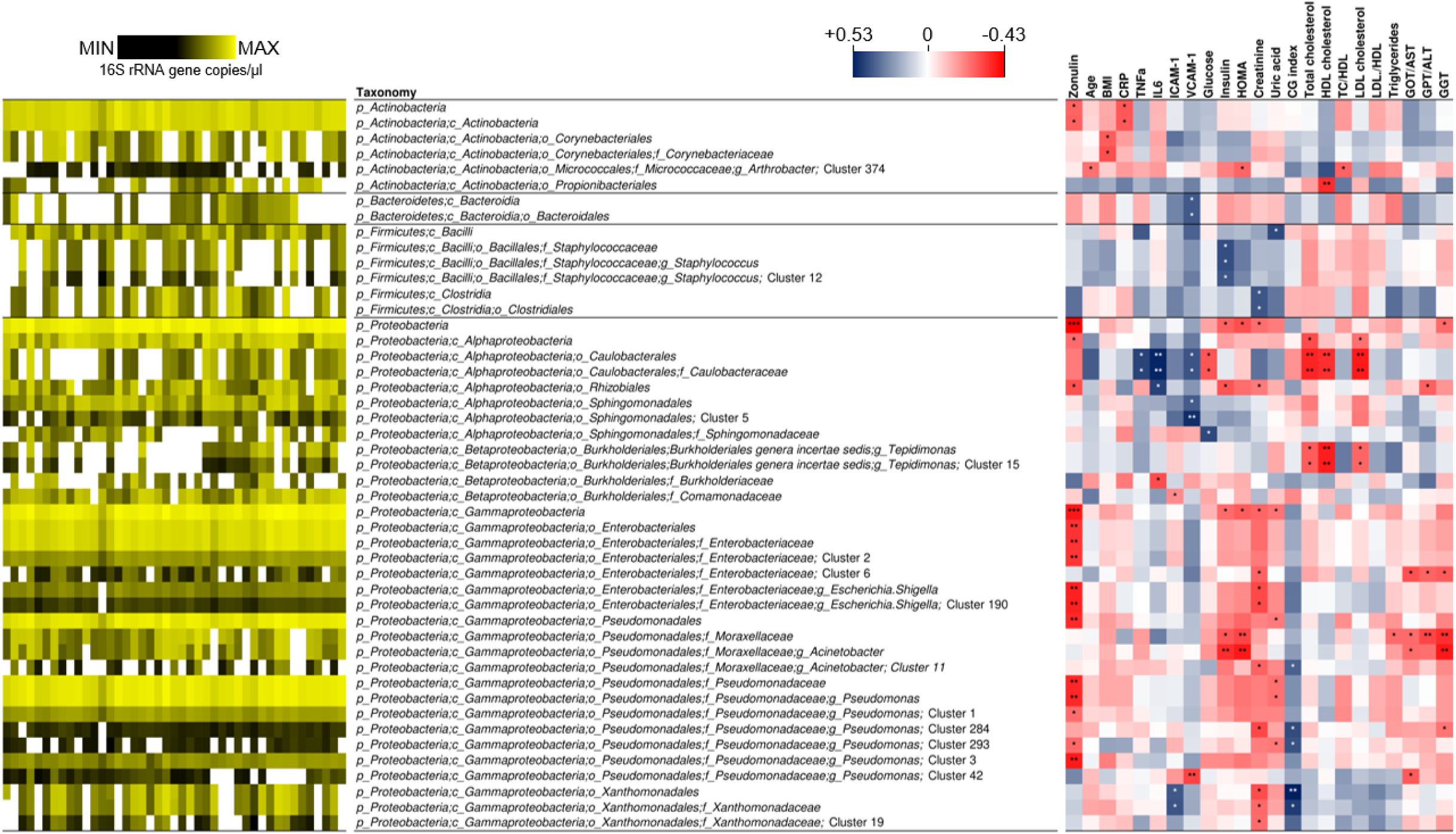
Correlations of the taxonomic units detected in blood toward age, BMI, and metabolic and functional markers determined in blood of the older subjects under study (n=43). This figure only includes taxa whose abundance significantly correlated with at least one parameter. The heatmap on the left (black-yellow color gradient) shows the distribution among samples of the abundances of taxonomic units. White boxes in the black-yellow heatmap indicate that taxa that have not been detected in a specific sample. The heatmap on the right (blue-white-red color gradient) represents the Spearman’s correlation coefficient, ρ. Asterisks indicate the P value of the Kendall’s rank correlation: *, P<0.05; **, P<0.01; ***, P<0.001.

Overall, these results indicate that serum zonulin levels are mostly associated with blood bacterial DNA ascribed to the class *γ-Proteobacteria* and to the genus *Pseudomonas*.

## Discussion

The main objective of this study was to quantify and characterize the bacterial DNA present in the bloodstream of older individuals, and to identify possible significant correlations with different markers relevant for health status in older people.

Our results demonstrate that quantifiable and highly variable amounts of bacterial DNA is present in the venous blood of older subjects. Reports of other studies that were concerned with healthy adult volunteers have also provided evidence of bacterial DNA in human blood, suggesting that it is a common phenomenon even in apparently healthy subjects (Nikkari et al 2001, Paisse et al 2016). Nonetheless, our study is the first to report this observation in an older population.

Based on the limited literature data available, the amount of bacterial DNA in blood appears to be highly variable between individual subjects when assessed by quantification of 16S rRNA gene copy number (Paisse et al 2016). For instance, the same protocol adopted in our study was previously used to quantify bacterial DNA in whole blood or buffy coat samples collected from healthy and obese subjects, and blood DNA concentrations ranging from zero up to about 8 × 10^7^ 16S rRNA gene copies per ml were obtained (Lelouvier et al 2016, Paisse et al 2016). In our study, we found an average of 5.8 (± 3.1) × 10^6^ copies of 16S rRNA gene per ml of whole blood (minimum of 1.3 × 10^6^ and maximum of 1.2 × 10^7^); therefore, the older volunteers studied here harbor levels of circulating bacterial DNA not dissimilar from those reported for other groups of individuals (Lelouvier et al 2016). Nonetheless, the data available to date are still too limited to draw conclusions.

The amount of bacterial DNA detected in blood was found to be significantly correlated with the serum level of zonulin. This significant correlation was confirmed when the analyses were repeated on a second set of blood samples, collected from the same older subjects after about 4 months, suggesting a stable association between these two variables in the group of subjects under investigation. To the best of our knowledge, the potential association between circulating bacterial DNA and zonulin has never been reported before. Zonulin has been proposed as a marker of intestinal permeability involved in chronic inflammatory conditions (Sturgeon and Fasano 2016), and has been shown to be increased in several pathologic states, including nonalcoholic fatty liver disease (Pacifico et al 2014), sepsis (Klaus et al 2013), obesity-associated insulin resistance (Moreno-Navarrete et al 2012), and diabetes (Jayashree et al 2014, Sapone et al 2006). Zonulin was also observed to be elevated in healthy aging (Qi et al 2017), supporting the hypothesis (recently verified in mouse model (Thevaranjan et al 2017)), that intestinal permeability increases with age, promoting the progressive entry into the bloodstream of microbial factors which trigger the immune system leading to the low-grade systemic inflammation typically associated with aging, Therefore, inflammaging may promote chronic-degenerative diseases (Franceschi and Campisi 2014). This mechanistic explanation for the onset of age-associated diseases is in agreement with the significant direct correlation observed between zonulin and the abundance of blood bacterial DNA in our study. Nonetheless, the putatively causal relationship between zonulin-mediated epithelial/endothelial permeability and bacterial DNAemia may be bidirectional because, whereas on one hand an increase in zonulin can lead to an increase in permeability and therefore to a greater translocation of bacterial DNA into the blood, on the other hand, the greater bacterial translocation in the blood can increase the local and systemic inflammation, which has been shown to induce zonulin expression (Moreno-Navarrete et al 2012).

In the second part of this study, we taxonomically characterized the 16S rRNA genes detected in blood samples. Most of the bacterial DNA detected in blood in our study was ascribed to the phylum *Proteobacteria* and *Actinobacteria*, whereas *Firmicutes* and *Bacteroidetes* were less represented and not detected in all samples. A similar distribution of phyla was described by Paisse et al., who analyzed buffy coat, plasma and red blood cells in a cohort of thirty healthy blood donors (Paisse et al 2016), and by Lelouvier et al., who studied a cohort of thirty-seven obese people (Lelouvier et al 2016). The dominance of *Proteobacteria* and, secondarily, *Actinobacteria* was also reported by other independent studies, in which the authors investigated groups of individuals with different health conditions such as acute pancreatitis and cardiovascular disease, as well as healthy controls (Li et al 2018, Nikkari et al 2001, Rajendhran et al 2013). A high proportion of *Proteobacteria* was also detected in the DNA isolated from murine muscle, liver and brain tissues (Lluch et al 2015), whereas in contrast, *Bacteroidetes* were reported to dominate the bacterial DNA detected in the blood of cats (Vientos-Plotts et al 2017), suggesting that the characteristics of DNAemia may be host-specific.

In a recent publication, circulating cell-free DNA isolated from human blood plasma was subjected to massive shotgun sequencing (Kowarsky et al 2017); more than half of the identified contigs had little or no homology with sequences in available databases and, interestingly, were assigned to hundreds of entirely novel microbial taxa. In our study, we did not find such a large presence of unknown microorganisms. Nonetheless, two main aspects distinguish the research by Kowarsky et al. from ours: (i) we performed 16S rRNA gene profiling and not shotgun metagenomic sequencing and (ii) we analyzed DNA isolated from whole blood and not plasma. This second aspect is particularly important considering the presence of bacterial DNA in blood cells such as erythrocytes and antigen-presenting cells (Minasyan 2016, Potgieter et al 2015).

The third most abundant genus, detected in all but one of the samples, was the *Actinobacteria* taxon *Arthrobacter*. Interestingly, members of this genus have been frequently isolated as viable cells from wounds and blood (see table 1 in (Mages et al 2008) which reported about 20 different *Arthrobacter* strains isolated from human blood and wound samples).

It is generally assumed that blood bacterial atopobiosis principally depends on translocation from the intestine (Lelouvier et al 2016); nonetheless, also the oral microbiota was suggested as a primary source of bacterial DNA in blood (Koren et al 2011, Ling et al 2017). Interestingly, in the study by Vientos-Plotts on cats, the taxonomic composition of blood was found to resemble the lung microbiota, which was determined analyzing bronchoalveolar lavage fluids (Vientos-Plotts et al 2017). In our study, some of the most represented bacteria found in blood samples belong to taxa that have been reported as dominant members of the lung microbiota, such as *Pseudomonas, Sphyngomonadales* and *Acinetobacter* (Beck et al 2012, Erb-Downward et al 2011, Zakharkina et al 2013). In addition, among the most represented taxa detected in blood, we also found members of the families *Comamonadaceae, Sphingomonadaceae*, and *Oxalobacteraceae*, which have been correlated with bronchial hypersensitivity in asthmatic patients (Huang et al 2011). These observations suggest that lung microbiota may be an additional source of bacterial DNA in blood. This hypothesis is not in contrast with our finding of a correlation between zonulin and blood bacterial DNA. In fact, zonulin, which is an emerging marker of intestinal permeability (Fasano 2012), was recently demonstrated to also be involved in the regulation of permeability in the lungs (Rittirsch et al 2013). More generally, although the origin of the bacterial DNA detected in blood remains substantially unknown, the evident association with zonulin serum levels suggests that bacteria may have translocated into the blood through a para-cellular route at epithelial and/or endothelial cell surfaces, an event that plausibly occurred not only in the intestine.

The blood concentration of DNA ascribed to several bacterial taxa significantly correlated with a few host metabolic factors. In particular, we found a significant correlation of *γ-Proteobacteria* and *Pseudomonas* with serum zonulin for both sets of blood samples (i.e., blood collected at recruitment and after approximately four months). However, the actual physiological meaning of such associations (if any) cannot be easily predicted and, more importantly, we believe that the taxonomic results reported here must be interpreted with caution for the reasons discussed below.

In this study, the bacterial DNA isolated from blood was taxonomically profiled through MiSeq sequencing of 16S rRNA gene amplicons. The use of this approach for the bacterial taxonomic characterization of low microbial biomass samples, such as blood, has been criticized as being at high risk of microbial contamination that may occur at any step of the protocol, from sample collection until sequencing (Eisenhofer et al 2019, Salter et al 2014, Weyrich et al 2019). In our study, we analyzed several control samples to assess the potential presence of contaminants in labware (e.g. vacutainer and EDTA tubes) and reagents (e.g. solutions used during extraction, library preparation, sequencing, and qPCR). According to qPCR experiments, we always detected in control samples a quantity of bacterial DNA much lower than that quantified in blood samples, suggesting the potential contaminants should not have significantly affected the taxonomic profiling of blood samples. Nonetheless, one limitation of this study is the absence of controls for the so-called “carrier effect”, i.e. the ability of additional nucleic acids (such as the host’s DNA and RNA that are abundantly present in blood) to enhance the recovery of DNA from low-abundant microbial cells in the sample (Xu et al 2009). However, the confirmation of a significant correlation between zonulin and16S rRNA gene copies in blood (total and ascribed to *Pseudomonas*) also in the second set of blood samples investigated supports the conclusion that the bacterial DNA detected in blood largely do not derive from contamination.

Considering the relative abundance of bacterial taxa detected in blood and control samples, we hypothesize that the most probable contaminants belong to the families *Enterobacteriaceae, Micrococcaceae* and *Moraxellaceae* (the second, third and fifth most abundant families detected in blood, respectively), whereas at least most part of the DNA ascribed to *Pseudomonadaceae* (the most abundant family detected in blood) is less likely to derive from contaminants. Lists of bacterial taxa that were identified in negative controls during different independent studies have been proposed (Eisenhofer et al 2019, Salter et al 2014), cataloging up to 70 different genera to be considered as potential contaminants (Eisenhofer et al 2019). These lists contain numerous *Proteobacteria* including *Pseudomonas*, which was found to be the most prevalent and abundant bacterial genus in the blood samples investigated in our study. *Pseudomonas* is a ubiquitous bacterium, which colonizes numerous environments, such as soil, water and various plant and animal organisms, due to minimal survival requirements and remarkable adaptation ability (Goldberg 2000). Notably, *Pseudomonas* is also one of the microorganisms most frequently isolated from patients with bacteremia, particularly the species *P. aeruginosa* (Nielsen 2015). In this report, the partial sequence of the 16S rRNA gene belonging to the most prevalent and abundant OTUs found in the analyzed blood samples (Cluster 1 and Cluster 3, Fig. 5) shared 100% similarity with *P. fluorescens* and other species of the same phylogenetic lineage. Although far less pathogenic than *P. aeruginosa, P. fluorescens* has been often reported as the aetiologic agent of opportunistic infections in lungs, mouth, stomach, urinary tract, skin, and, most commonly, blood (Azam and Khan 2019, Scales et al 2014). Notably, *P. fluorescens* is recognized as the most important cause of iatrogenic sepsis, attributed to contaminated blood transfusion or contaminated equipment used in intravenous infusions (Benito et al 2012, Centers for Disease and Prevention 2005, Centers for Disease and Prevention 2006).

Although the literature evidence discussed above suggests that *P. fluorescens* and related species can be contaminants, on the other hand, these bacteria were also reported to possess numerous functional properties that support their survival and growth in mammalian hosts (Scales et al 2014). Furthermore, an interesting association was found between the presence of serum antibodies against the I2 peptide encoded by *P. fluorescens* and Crohn’s disease (Landers et al 2002), celiac disease (Ashorn et al 2008), ankylosing spondylitis (Mundwiler et al 2009), and chronic granulomatous disease (Yu et al 2011). In addition, *P. fluorescens* was reported to be regularly cultured from clinical samples even in the absence of acute infection (Scales et al 2014). Finally, *P. fluorescens* was demonstrated to induce zonulin expression and decreased intestinal permeability in a time dependent manner in an in vitro model of intestinal epithelium. In the same study, the authors found increased zonulin levels and higher abundance of *Pseudomonas* 16S rRNA gene copies (as determined through qPCR with genus-specific primers) in coronary artery disease (CAD) patients compared to non-CAD subjects (Li et al 2016). Altogether, these reports support the hypothesis that human-adapted *P. fluorescens* strains constitute low-abundance indigenous members of the microbial ecosystem of various body sites, such as the lungs, mouth, and stomach (Diaz et al 2013, Dickson et al 2014, Patel et al 2013, Scales et al 2014). Contextually, we can speculate that certain *P. fluorescens*-related strains are highly adaptable and poorly pathogenic members of the microbiota in several body sites that may frequently translocate into the bloodstream, providing a dominant contribution to bacterial DNAemia. However, we are conscious that our results do not conclusively demonstrate the actual presence of *Pseudomonas* (cells or free DNA) in blood. We believe that DNA-independent methods, for instance based on the use of electron microscopy or bacterium-specific antibodies, could contribute to the unambiguous demonstration of the presence of these bacteria in human blood.

## Conclusions

Based on the data reported here and in the context of previous reports of DNAemia in younger adults, the authors conclude that older individuals harbor detectable amounts of bacterial DNA in their blood and that it has a somewhat distinct taxonomic origin. In addition, our observed correlation between the concentration of bacterial DNA in blood and the serum levels of zonulin, an emerging marker of intestinal permeability, suggest that DNAemia is directly related with paracellular permeability, including but not necessarily limited to the epithelium of the gastrointestinal tract. We speculate that the bacterial DNA detected in blood, irrespective of the origin (gut, oral cavity or lung) and the form (free DNA, free bacterial cells, or bacteria internalized in blood cells), is indicative of a bacterial DNAemia that is not-silent, but may influence several aspects of host physiology. In particular, we propose that bacterial DNAemia may represent an important candidate biomarker helpful for the prognosis and/or prediction of metabolic and clinical pathologic conditions in the older subjects and can also be actively implicated in the development of inflammaging and aging associated diseases.

## Acknowledgements

We warmly thank all the volunteers participating to the study for their valuable contribution. We are grateful to Alberto Fantuzzo, Chiara Cavazzini, Lorella Pinton, Paolo Bergantin, Rosanna Ceccato, Pamela Soranzo, and Silvana Giraldini at Opera Immacolata Concezione (OIC Foundation, Padua, Italy) for their coordinating activities in the nursing home. We are also grateful to all physicians (Michela Rigon, Lorena D’Aloise, Antonio Merlo, Elisabetta Bernardinello, Nadia Malacarne, Silvana Bortoli, Fabiola Talato, Agostino Corsini, Maria Licursi, Nicoletta Marcon, Angela Sansone), nurses and other personnel at OIC who were essential to complete the study successfully.

## Funding

The project coordinated by P. Riso and S. Guglielmetti “Gut and blood microbiomics for studying the effect of a polyphenol-rich dietary pattern on intestinal permeability in the elderly (MaPLE)” was funded under the Intestinal-Microbiomics call (2015) of the Joint Programming Initiative, “A Healthy Diet for a Healthy Life” (JPI HDHL, website: http://www.healthydietforhealthylife.eu). This project was funded by the respective national research councils: Mipaaf (Italy; D.M. 8245/7303/2016), the Biotechnology and Biological Sciences Research Council (BBSRC, UK; Grant BB/R012512/1) and the Ministry of Economy and Competitiveness (MINECO) (Spain; PCIN-PCIN-2015-238 MINECO). Additional funding was provided by the Generalitat de Catalunya’s Agency AGAUR Spain; grant no. 2017SGR1546), CIBERFES (co-funded by the FEDER Program from EU), and the BBSRC (UK) through an Institute Strategic Programme Grant (‘Food Innovation and Health’; Grant No. BB/R012512/1 and its constituent projects BBS/E/F/000PR10343, BBS/E/F/000PR10345 and BBS/E/F/000PR10346) to the Quadram Institute Bioscience.

## Availability of data and materials

The MaPLE trial was registered at the *International Standard Randomised Controlled Trial Number* (ISRCTN) registry under the code ISRCTN10214981. According to the data management plan of the MaPLE project, the datasets or analyzed during the current study are available in the Dataverse repository, https://dataverse.unimi.it/dataverse/JPI-MaPLE (dataset named “Blood Microbiomics data at baseline”). Blood microbiomics sequencing reads have been also deposited in the European Nucleotide Archive (ENA) of the European Bioinformatics Institute under accession code PRJEB30560.

## Authors’ contributions

PR, and SG conceived and designed the study and equally contributed to the work. AC, CAL and PK contributed significantly to the development of the experimental protocol. AC supervised older subjects’ enrollment both considering inclusion criteria and clinical parameters evaluation. PR, CDB and SB recruited volunteers, collected samples, and performed zonulin measurements and blood biochemistry analyses. MSW and PAK performed the analyses of immunological parameters. GG, SG and VT performed the bioinformatic analysis of microbiomic data and the statistical analyses. SG wrote the manuscript with PR support. All authors critically reviewed the manuscript.

## Ethics approval

The study was approved by the Research Ethics Committee of the Università degli Studi di Milano (opinion no. 6/16; February 15^th^, 2016).

## Competing interests

The authors have no conflict of interest to declare.

## Additional file 1

The document contains Supplementary Figures S1-S5.

## Notes

https://dataverse.unimi.it/dataverse/JPI-MaPLE

### Competing Interest Statement

The authors have declared no competing interest.

## References

Amar J, Serino M, Lange C, Chabo C, Iacovoni J, Mondot S et al (2011). Involvement of tissue bacteria in the onset of diabetes in humans: evidence for a concept. Diabetologia 54: 3055–3061.

Amar J, Lange C, Payros G, Garret C, Chabo C, Lantieri O et al (2013). Blood microbiota dysbiosis is associated with the onset of cardiovascular events in a large general population: the D.E.S.I.R. study. PloS one 8: e54461.

Ashorn S, Raukola H, Valineva T, Ashorn M, Wei B, Braun J et al (2008). Elevated serum anti-Saccharomyces cerevisiae, anti-I2 and anti-OmpW antibody levels in patients with suspicion of celiac disease. Journal of clinical immunology 28: 486–494.

Azam MW, Khan AU (2019). Updates on the pathogenicity status of Pseudomonas aeruginosa. Drug discovery today 24: 350–359.

Beck JM, Young VB, Huffnagle GB (2012). The microbiome of the lung. Transl Res 160: 258–266.

Benito N, Mirelis B, Luz Galvez M, Vila M, Lopez-Contreras J, Cotura A et al (2012). Outbreak of Pseudomonas fluorescens bloodstream infection in a coronary care unit. The Journal of hospital infection 82: 286–289.

Centers for Disease C, Prevention (2005). Pseudomonas bloodstream infections associated with a heparin/saline flush--Missouri, New York, Texas, and Michigan, 2004-2005. MMWR Morbidity and mortality weekly report 54: 269–272.

Centers for Disease C, Prevention (2006). Update: Delayed onset Pseudomonas fluorescens bloodstream infections after exposure to contaminated heparin flush--Michigan and South Dakota, 2005-2006. MMWR Morbidity and mortality weekly report 55: 961–963.

Collado MC, Rautava S, Aakko J, Isolauri E, Salminen S (2016). Human gut colonisation may be initiated in utero by distinct microbial communities in the placenta and amniotic fluid. Sci Rep 6: 23129.

Diaz PI, Hong BY, Frias-Lopez J, Dupuy AK, Angeloni M, Abusleme L et al (2013). Transplantation-associated long-term immunosuppression promotes oral colonization by potentially opportunistic pathogens without impacting other members of the salivary bacteriome. Clinical and vaccine immunology: CVI 20: 920–930.

Dickson RP, Erb-Downward JR, Freeman CM, Walker N, Scales BS, Beck JM et al (2014). Changes in the lung microbiome following lung transplantation include the emergence of two distinct Pseudomonas species with distinct clinical associations. PloS one 9: e97214.

Eisenhofer R, Minich JJ, Marotz C, Cooper A, Knight R, Weyrich LS (2019). Contamination in Low Microbial Biomass Microbiome Studies: Issues and Recommendations. Trends in microbiology 27: 105–117.

Erb-Downward JR, Thompson DL, Han MK, Freeman CM, McCloskey L, Schmidt LA et al (2011). Analysis of the lung microbiome in the “healthy” smoker and in COPD. PloS one 6: e16384.

Escudie F, Auer L, Bernard M, Mariadassou M, Cauquil L, Vidal K et al (2018). FROGS: Find, Rapidly, OTUs with Galaxy Solution. Bioinformatics 34: 1287–1294.

Fasano A (2012). Intestinal permeability and its regulation by zonulin: diagnostic and therapeutic implications. Clin Gastroenterol Hepatol 10: 1096–1100.

Franceschi C, Bonafe M, Valensin S, Olivieri F, De Luca M, Ottaviani E et al (2000). Inflamm-aging. An evolutionary perspective on immunosenescence. Ann N Y Acad Sci 908: 244–254.

Franceschi C, Campisi J (2014). Chronic inflammation (inflammaging) and its potential contribution to age-associated diseases. J Gerontol A Biol Sci Med Sci 69 Suppl 1: S4–9.

Gargari G, Deon V, Taverniti V, Gardana C, Denina M, Riso P et al (2018a). Evidence of dysbiosis in the intestinal microbial ecosystem of children and adolescents with primary hyperlipidemia and the potential role of regular hazelnut intake. FEMS microbiology ecology 94.

Gargari G, Taverniti V, Gardana C, Cremon C, Canducci F, Pagano I et al (2018b). Fecal Clostridiales distribution and short-chain fatty acids reflect bowel habits in irritable bowel syndrome. Environmental microbiology 20: 3201–3213.

Goldberg JB (2000). Pseudomonas: global bacteria. Trends in microbiology 8: 55–57.

Gosiewski T, Ludwig-Galezowska AH, Huminska K, Sroka-Oleksiak A, Radkowski P, Salamon D et al (2017). Comprehensive detection and identification of bacterial DNA in the blood of patients with sepsis and healthy volunteers using next-generation sequencing method - the observation of DNAemia. Eur J Clin Microbiol Infect Dis 36: 329–336.

Guglielmetti S, Bernardi S, Del Bo C, Cherubini A, Porrini M, Gargari G et al (2020). Effect of a polyphenol-rich dietary pattern on intestinal permeability and gut and blood microbiomics in older subjects: study protocol of the MaPLE randomised controlled trial. BMC geriatrics 20: 77.

Huang YJ, Nelson CE, Brodie EL, Desantis TZ, Baek MS, Liu J et al (2011). Airway microbiota and bronchial hyperresponsiveness in patients with suboptimally controlled asthma. J Allergy Clin Immunol 127: 372-381 e371-373.

Jayashree B, Bibin YS, Prabhu D, Shanthirani CS, Gokulakrishnan K, Lakshmi BS et al (2014). Increased circulatory levels of lipopolysaccharide (LPS) and zonulin signify novel biomarkers of proinflammation in patients with type 2 diabetes. Mol Cell Biochem 388: 203–210.

Klaus DA, Motal MC, Burger-Klepp U, Marschalek C, Schmidt EM, Lebherz-Eichinger D et al (2013). Increased plasma zonulin in patients with sepsis. Biochem Med (Zagreb) 23: 107–111.

Koren O, Spor A, Felin J, Fak F, Stombaugh J, Tremaroli V et al (2011). Human oral, gut, and plaque microbiota in patients with atherosclerosis. Proceedings of the National Academy of Sciences of the United States of America 108 Suppl 1: 4592–4598.

Kowarsky M, Camunas-Soler J, Kertesz M, De Vlaminck I, Koh W, Pan W et al (2017). Numerous uncharacterized and highly divergent microbes which colonize humans are revealed by circulating cell-free DNA. Proceedings of the National Academy of Sciences of the United States of America 114: 9623–9628.

Landers CJ, Cohavy O, Misra R, Yang H, Lin YC, Braun J et al (2002). Selected loss of tolerance evidenced by Crohn’s disease-associated immune responses to auto- and microbial antigens. Gastroenterology 123: 689–699.

Lelouvier B, Servant F, Paisse S, Brunet AC, Benyahya S, Serino M et al (2016). Changes in blood microbiota profiles associated with liver fibrosis in obese patients: A pilot analysis. Hepatology 64: 2015–2027.

Li C, Gao M, Zhang W, Chen C, Zhou F, Hu Z et al (2016). Zonulin Regulates Intestinal Permeability and Facilitates Enteric Bacteria Permeation in Coronary Artery Disease. Sci Rep 6: 29142.

Li Q, Wang C, Tang C, Zhao X, He Q, Li J (2018). Identification and Characterization of Blood and Neutrophil-Associated Microbiomes in Patients with Severe Acute Pancreatitis Using Next-Generation Sequencing. Front Cell Infect Microbiol 8: 5.

Ling Z, Liu X, Cheng Y, Shao L, Jiang H, Li L (2017). Blood microbiota as a potential noninvasive diagnostic biomarker for liver fibrosis in severely obese patients: Choose carefully. Hepatology 65: 1775–1776.

Lluch J, Servant F, Paisse S, Valle C, Valiere S, Kuchly C et al (2015). The Characterization of Novel Tissue Microbiota Using an Optimized 16S Metagenomic Sequencing Pipeline. PloS one 10: e0142334.

Ma TY, Hollander D, Dadufalza V, Krugliak P (1992). Effect of aging and caloric restriction on intestinal permeability. Exp Gerontol 27: 321–333.

Mages IS, Frodl R, Bernard KA, Funke G (2008). Identities of Arthrobacter spp. and Arthrobacter-like bacteria encountered in human clinical specimens. J Clin Microbiol 46: 2980–2986.

Metchnikoff E (1907). The prolongation of life: optimistic studies. William Heinemann: London

Minasyan H (2016). Mechanisms and pathways for the clearance of bacteria from blood circulation in health and disease. Pathophysiology: the official journal of the International Society for Pathophysiology 23: 61–66.

Moreno-Navarrete JM, Sabater M, Ortega F, Ricart W, Fernandez-Real JM (2012). Circulating zonulin, a marker of intestinal permeability, is increased in association with obesity-associated insulin resistance. PloS one 7: e37160.

Mundwiler ML, Mei L, Landers CJ, Reveille JD, Targan S, Weisman MH (2009). Inflammatory bowel disease serologies in ankylosing spondylitis patients: a pilot study. Arthritis research & therapy 11: R177.

Nadkarni MA, Martin FE, Jacques NA, Hunter N (2002). Determination of bacterial load by real-time PCR using a broad-range (universal) probe and primers set. Microbiology 148: 257–266.

Nielsen SL (2015). The incidence and prognosis of patients with bacteremia. Danish medical journal 62.

Nikkari S, McLaughlin IJ, Bi W, Dodge DE, Relman DA (2001). Does blood of healthy subjects contain bacterial ribosomal DNA? J Clin Microbiol 39: 1956–1959.

Pacifico L, Bonci E, Marandola L, Romaggioli S, Bascetta S, Chiesa C (2014). Increased circulating zonulin in children with biopsy-proven nonalcoholic fatty liver disease. World journal of gastroenterology 20: 17107–17114.

Paisse S, Valle C, Servant F, Courtney M, Burcelin R, Amar J et al (2016). Comprehensive description of blood microbiome from healthy donors assessed by 16S targeted metagenomic sequencing. Transfusion 56: 1138–1147.

Patel SK, Pratap CB, Verma AK, Jain AK, Dixit VK, Nath G (2013). Pseudomonas fluorescens-like bacteria from the stomach: a microbiological and molecular study. World journal of gastroenterology 19: 1056–1067.

Potgieter M, Bester J, Kell DB, Pretorius E (2015). The dormant blood microbiome in chronic, inflammatory diseases. FEMS Microbiol Rev 39: 567–591.

Qi Y, Goel R, Kim S, Richards EM, Carter CS, Pepine CJ et al (2017). Intestinal Permeability Biomarker Zonulin is Elevated in Healthy Aging. J Am Med Dir Assoc 18: 810 e811–810 e814.

Rajendhran J, Shankar M, Dinakaran V, Rathinavel A, Gunasekaran P (2013). Contrasting circulating microbiome in cardiovascular disease patients and healthy individuals. Int J Cardiol 168: 5118–5120.

Rittirsch D, Flierl MA, Nadeau BA, Day DE, Huber-Lang MS, Grailer JJ et al (2013). Zonulin as prehaptoglobin2 regulates lung permeability and activates the complement system. Am J Physiol Lung Cell Mol Physiol 304: L863–872.

Salter SJ, Cox MJ, Turek EM, Calus ST, Cookson WO, Moffatt MF et al (2014). Reagent and laboratory contamination can critically impact sequence-based microbiome analyses. BMC biology 12: 87.

Sapone A, de Magistris L, Pietzak M, Clemente MG, Tripathi A, Cucca F et al (2006). Zonulin upregulation is associated with increased gut permeability in subjects with type 1 diabetes and their relatives. Diabetes 55: 1443–1449.

Sato J, Kanazawa A, Ikeda F, Yoshihara T, Goto H, Abe H et al (2014). Gut dysbiosis and detection of “live gut bacteria” in blood of Japanese patients with type 2 diabetes. Diabetes Care 37: 2343–2350.

Scales BS, Dickson RP, LiPuma JJ, Huffnagle GB (2014). Microbiology, genomics, and clinical significance of the Pseudomonas fluorescens species complex, an unappreciated colonizer of humans. Clinical microbiology reviews 27: 927–948.

Sturgeon C, Fasano A (2016). Zonulin, a regulator of epithelial and endothelial barrier functions, and its involvement in chronic inflammatory diseases. Tissue Barriers 4: e1251384.

Thevaranjan N, Puchta A, Schulz C, Naidoo A, Szamosi JC, Verschoor CP et al (2017). Age-Associated Microbial Dysbiosis Promotes Intestinal Permeability, Systemic Inflammation, and Macrophage Dysfunction. Cell Host Microbe 21: 455–466 e454.

Thomas-White K, Brady M, Wolfe AJ, Mueller ER (2016). The bladder is not sterile: History and current discoveries on the urinary microbiome. Curr Bladder Dysfunct Rep 11: 18–24.

Vientos-Plotts AI, Ericsson AC, Rindt H, Grobman ME, Graham A, Bishop K et al (2017). Dynamic changes of the respiratory microbiota and its relationship to fecal and blood microbiota in healthy young cats. PloS one 12: e0173818.

von Neubeck M, Huptas C, Gluck C, Krewinkel M, Stoeckel M, Stressler T et al (2017). Pseudomonas lactis sp. nov. and Pseudomonas paralactis sp. nov., isolated from bovine raw milk. International journal of systematic and evolutionary microbiology 67: 1656–1664.

Weyrich LS, Farrer AG, Eisenhofer R, Arriola LA, Young J, Selway CA et al (2019). Laboratory contamination over time during low-biomass sample analysis. Molecular ecology resources 19: 982–996.

Xu Z, Zhang F, Xu B, Tan J, Li S, Jin L (2009). Improving the sensitivity of negative controls in ancient DNA extractions. Electrophoresis 30: 1282–1285.

Yu JE, De Ravin SS, Uzel G, Landers C, Targan S, Malech HL et al (2011). High levels of Crohn’s disease-associated anti-microbial antibodies are present and independent of colitis in chronic granulomatous disease. Clinical immunology 138: 14–22.

Zakharkina T, Heinzel E, Koczulla RA, Greulich T, Rentz K, Pauling JK et al (2013). Analysis of the airway microbiota of healthy individuals and patients with chronic obstructive pulmonary disease by T-RFLP and clone sequencing. PloS one 8: e68302.

