## Supplemental material for "Bacterial DNAemia is associated with serum zonulin levels in older subjects"

**Fig. S1.** Linear regression and correlation analysis carried out between blood bacterial load and serum zonulin levels. These analyses were performed on the second set of blood samples (i.e., collected approximately four months after the first draw). Correlation has been assessed through Pearson's (r coefficient), Spearman's (ρ), and Kendall's (τ) analyses.

| | Pearson's | Spearman's | Kendall's |
|---|---|---|---|
| *r*, *ρ*, *τ* | 0.407 | 0.449 | 0.309 |
| p value | 0.0075 | 0.0029 | 0.0039 |

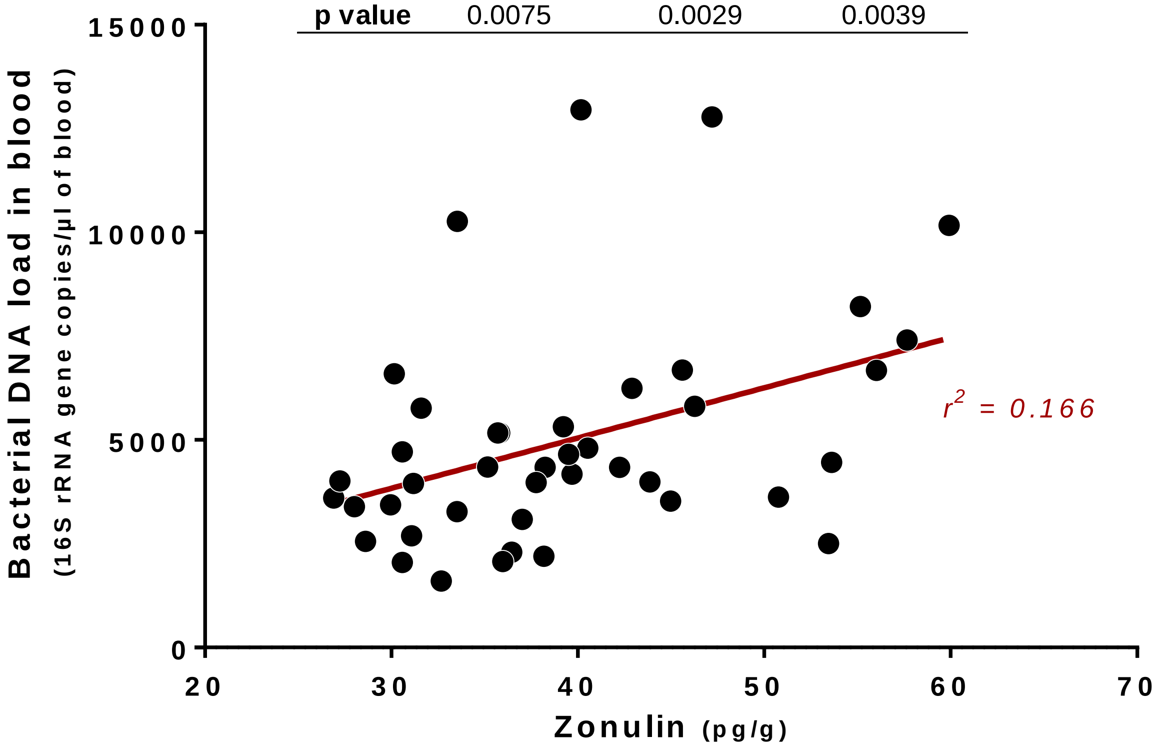

**Fig. S2**. Diversity analyses of data generated by the 16S rRNA gene profiling of blood bacterial samples and $C_{qPCR}$ controls. **A**, rarefaction curves based on the richness of operative taxonomic units (OTUs). **B**, α-diversity analysis based on five different indexes, shown in order of increasing evenness weight in the algorithm, from observed species and Chao1 (no evenness considered) to InvSimpson index. **C**, principal coordinates analysis of generalized UniFrac distance; the first two coordinates are displayed; the percentage of variance explained is indicated in brackets.

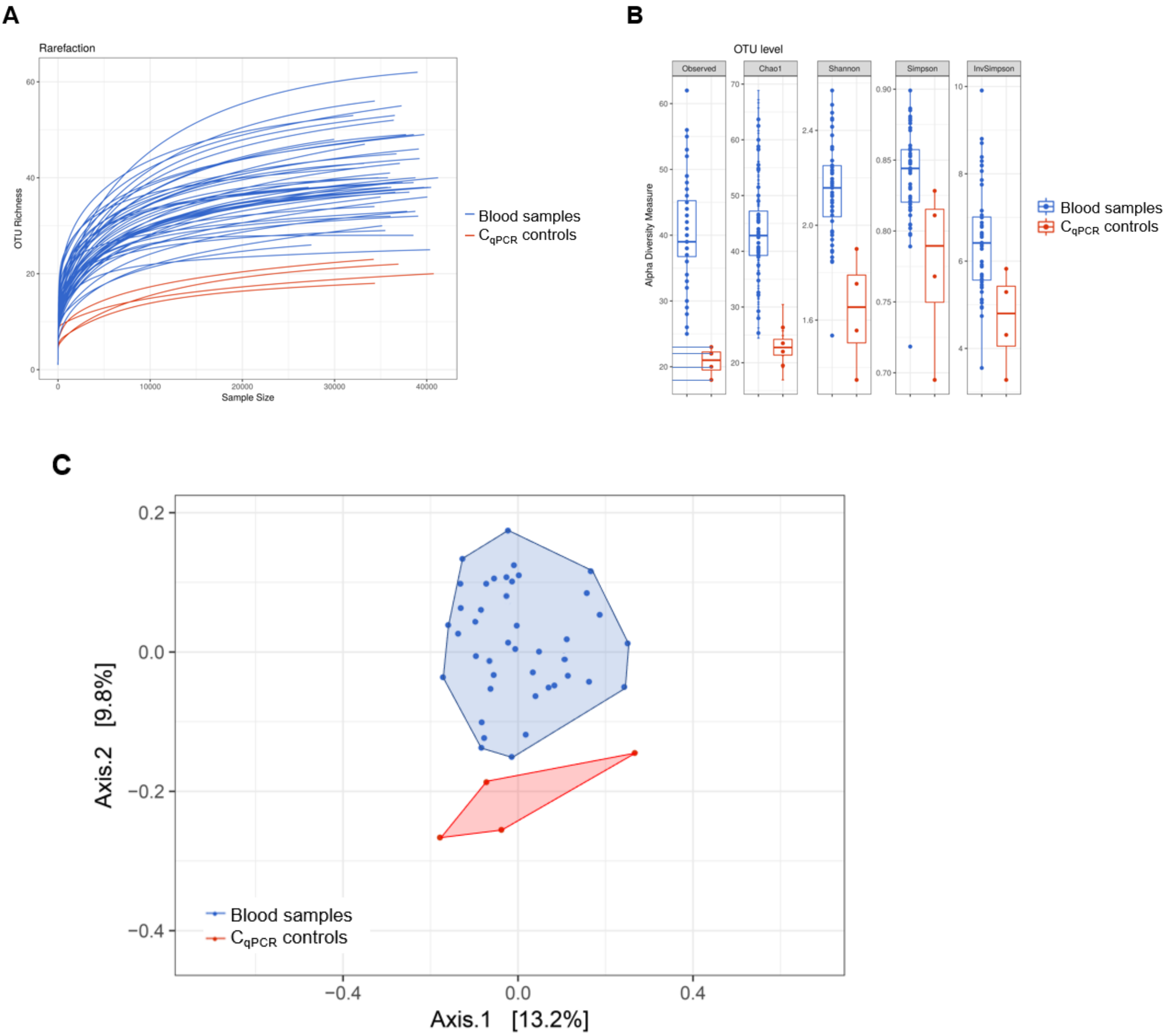

**Fig. S3**. Abundance in blood samples (B.S.) and controls ($C_{qPCR}$) of the main families (panels **A** and **C**) and genera (panels **B** and **D**) detected in blood (taxa are according to Fig. 4). Panels **A** and **B**, normalized abundances obtained multiplying the total 16S rRNA gene copies/µl by the percentage of each taxon in a specific sample. Panels **C** and **D**, relative abundances shown as percentage.

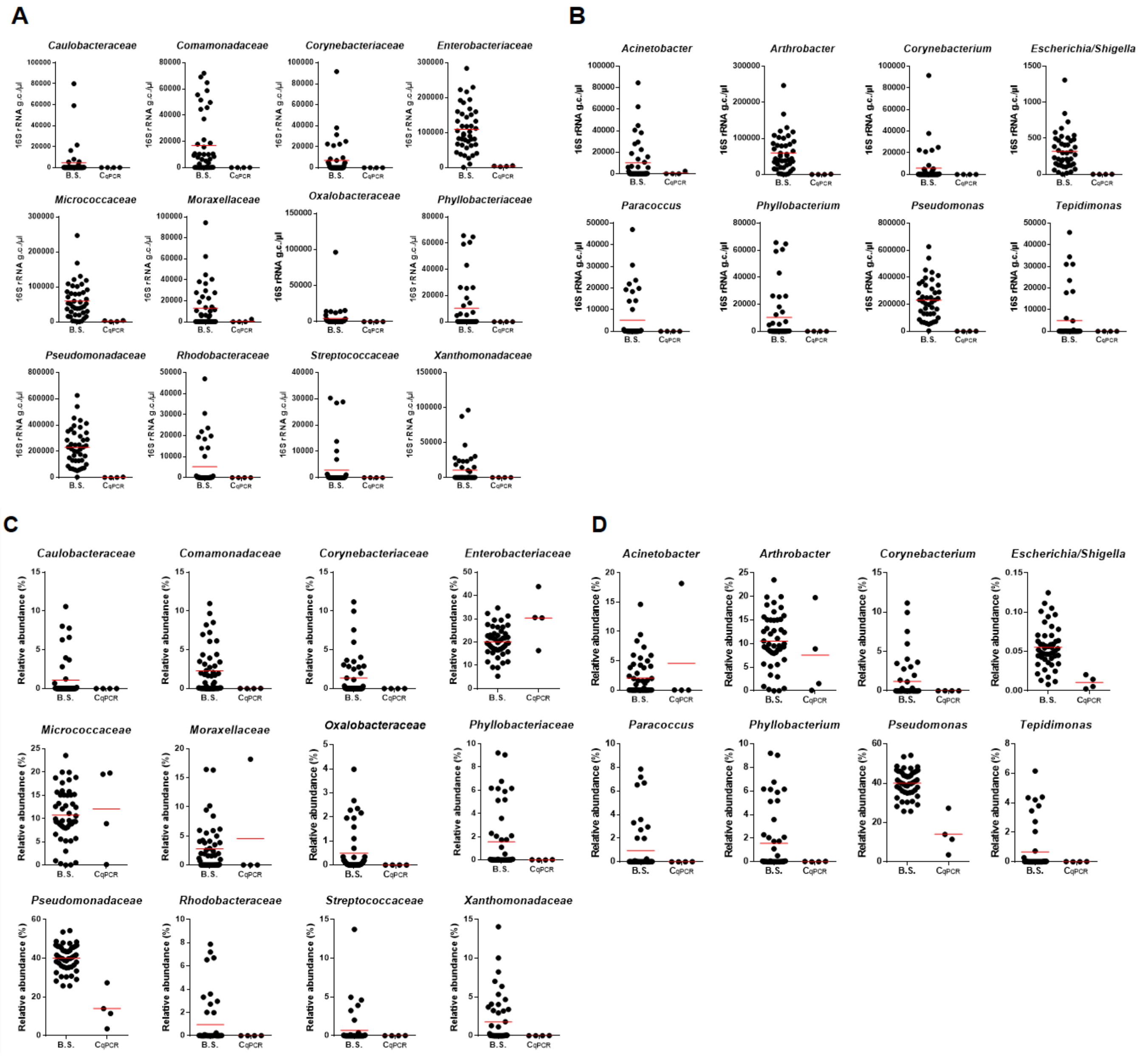

**Fig. S4**. Bacterial taxonomic profiling by 16S rRNA gene sequencing of additional controls and a blood sample (B.S.). $C_{extr}$, DNA extracted from ultrapure water; $C_{dEDTA}$, DNA extracted from commercial PBS; $C_{dEDTA}$, DNA extracted from commercial PBS passed through the vacutainer system. DNA extraction from all controls have been carried out following the same protocol used for the extraction of DNA from blood and was performed contemporaneously to blood sample B.S. **A**, concentration of 16S rRNA gene copies (16S rRNA g.c./μl) determined by qPCR. **B**, number of sequencing reads assigned to operational taxonomic units (OTUs) per sample. **C**, taxonomic composition of each sample at OTU level; each Cluster corresponds to an OTU; genera of main Clusters detected in B.S. are specified on the right. **D**, distribution of taxonomic Clusters (i.e., OTUs) detected in the analyzed samples; the bacterial genus to which each cluster belong is shown on the right; white cells in the heatmap indicate that the bacterial taxon was not detected in a specific sample.

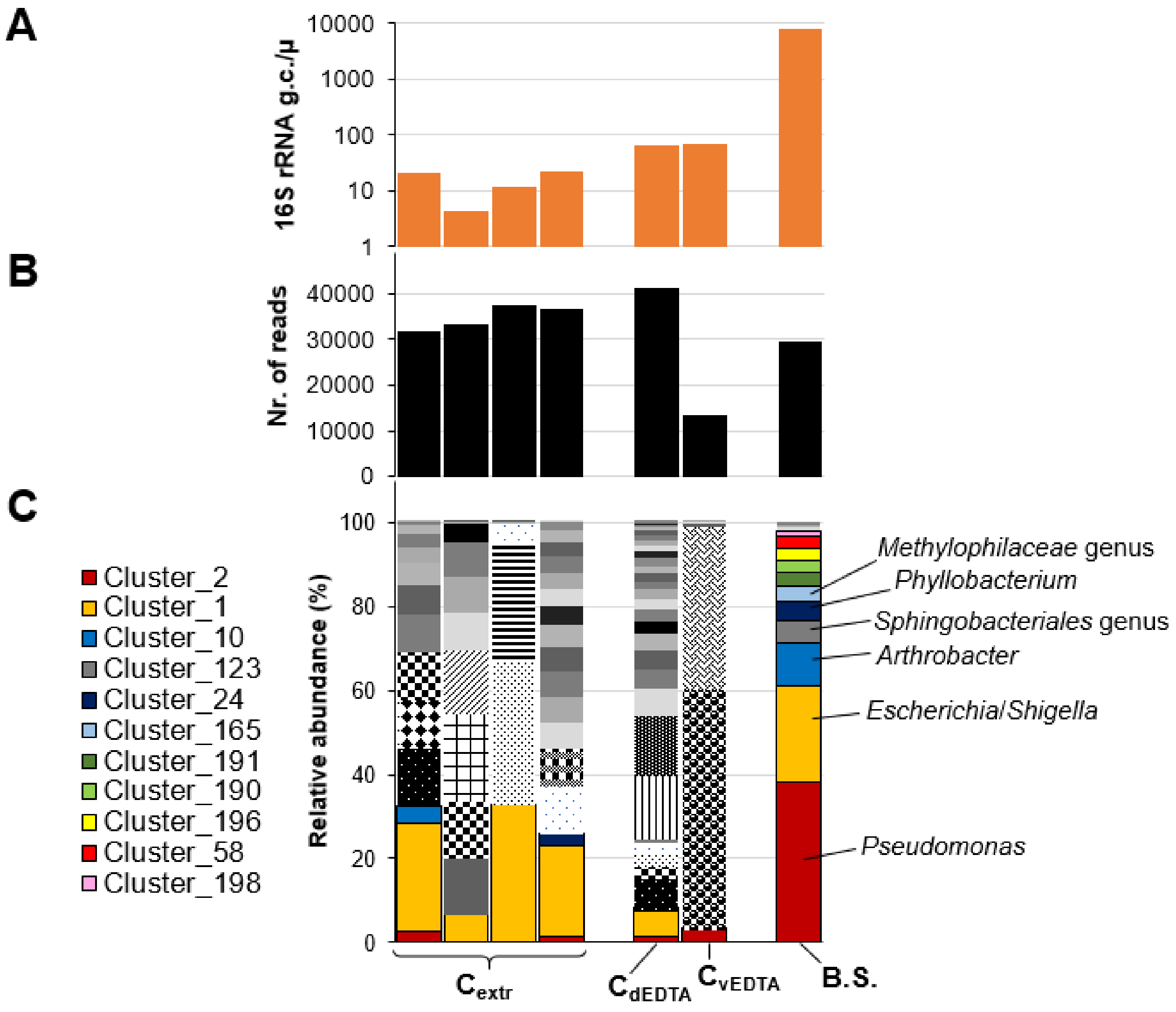

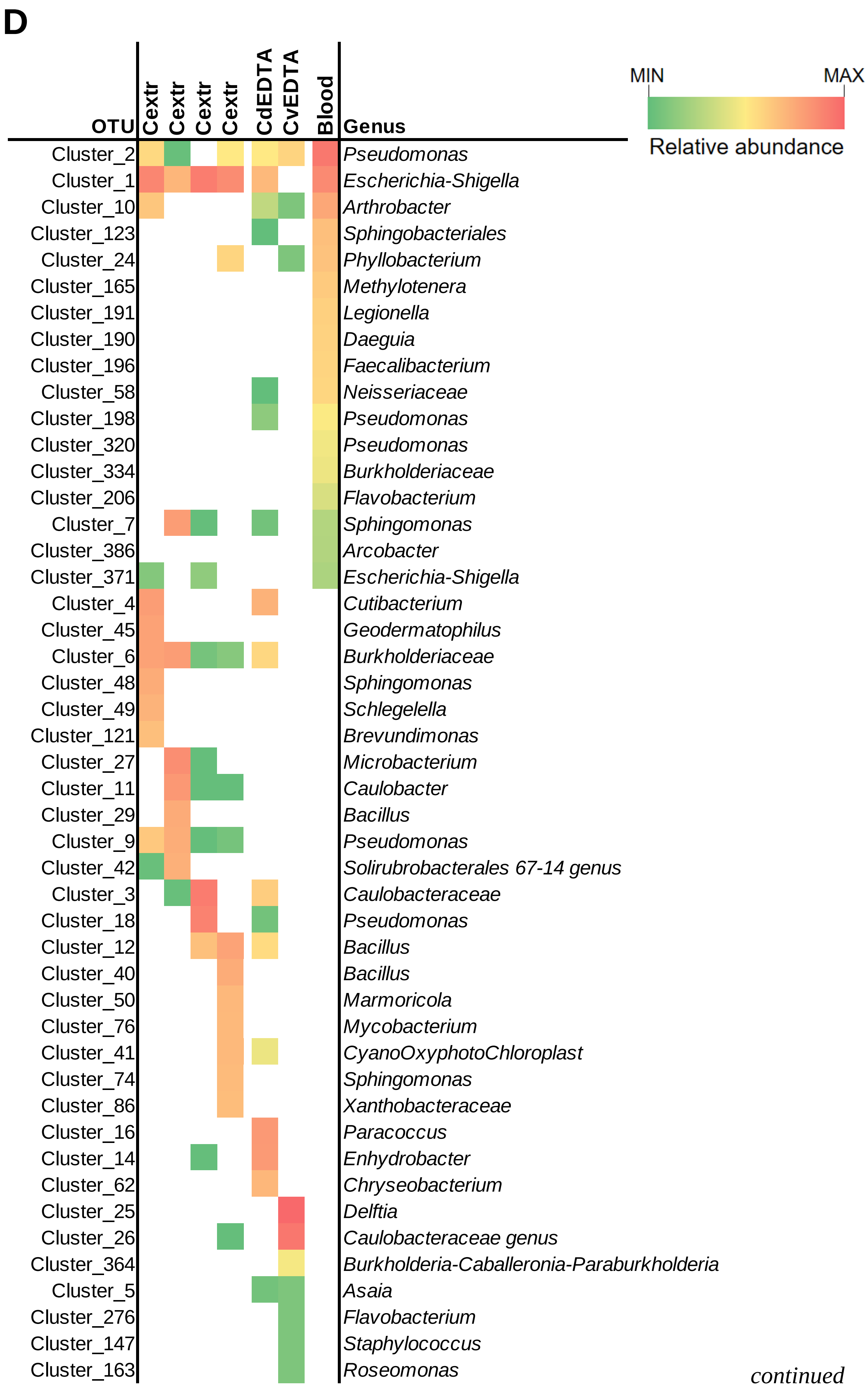
D
OTU
Cextr
Cextr
Cextr
Cextr
CdEDTA
CvEDTA
Blood
Genus
MIN
MAX
Relative abundance
Cluster_2 Pseudomonas
Cluster_1 Escherichia-Shigella
Cluster_10 Arthrobacter
Cluster_123 Sphingobacteriales
Cluster_24 Phyllobacterium
Cluster_165 Methylotenera
Cluster_191 Legionella
Cluster_190 Daeguia
Cluster_196 Faecalibacterium
Cluster_58 Neisseriaceae
Cluster_198 Pseudomonas
Cluster_320 Pseudomonas
Cluster_334 Burkholderiaceae
Cluster_206 Flavobacterium
Cluster_7 Sphingomonas
Cluster_386 Arcobacter
Cluster_371 Escherichia-Shigella
Cluster_4 Cutibacterium
Cluster_45 Geodermatophilus
Cluster_6 Burkholderiaceae
Cluster_48 Sphingomonas
Cluster_49 Schlegelella
Cluster_121 Brevundimonas
Cluster_27 Microbacterium
Cluster_11 Caulobacter
Cluster_29 Bacillus
Cluster_9 Pseudomonas
Cluster_42 Solirubrobacterales 67-14 genus
Cluster_3 Caulobacteraceae
Cluster_18 Pseudomonas
Cluster_12 Bacillus
Cluster_40 Bacillus
Cluster_50 Marmoricola
Cluster_76 Mycobacterium
Cluster_41 CyanoOxyphotoChloroplast
Cluster_74 Sphingomonas
Cluster_86 Xanthobacteraceae
Cluster_16 Paracoccus
Cluster_14 Enhydrobacter
Cluster_62 Chryseobacterium
Cluster_25 Delftia
Cluster_26 Caulobacteraceae genus
Cluster_364 Burkholderia-Caballeronia-Paraburkholderia
Cluster_5 Asaia
Cluster_276 Flavobacterium
Cluster_147 Staphylococcus
Cluster_163 Roseomonas

continued

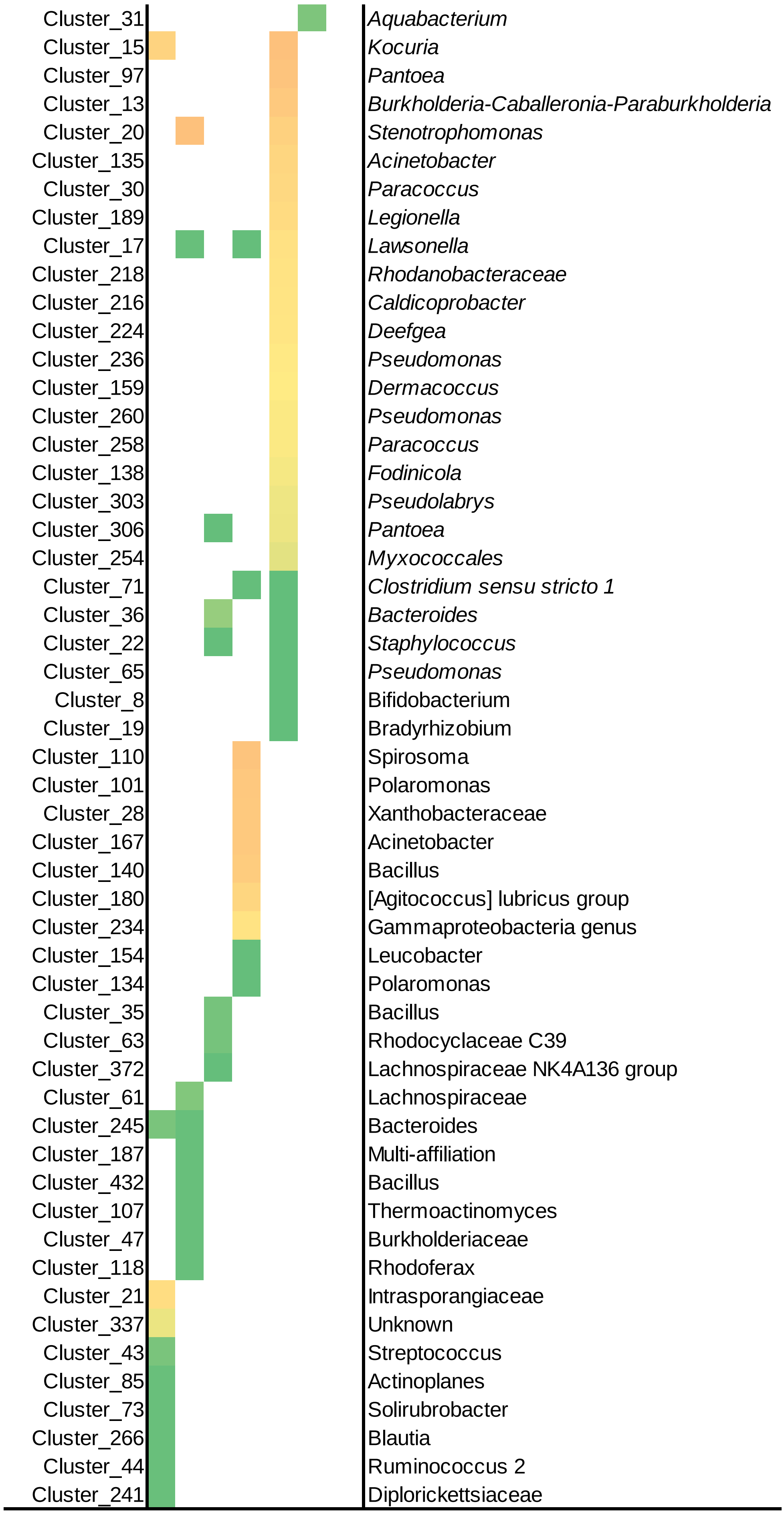
Cluster_31 Aquabacterium
Cluster_15 Kocuria
Cluster_97 Pantoea
Cluster_13 Burkholderia-Caballeronia-Paraburkholderia
Cluster_20 Stenotrophomonas
Cluster_135 Acinetobacter
Cluster_30 Paracoccus
Cluster_189 Legionella
Cluster_17 Lawsonella
Cluster_218 Rhodanobacteraceae
Cluster_216 Caldicoprobacter
Cluster_224 Deefgea
Cluster_236 Pseudomonas
Cluster_159 Dermacoccus
Cluster_260 Pseudomonas
Cluster_258 Paracoccus
Cluster_138 Fodinicola
Cluster_303 Pseudolabrys
Cluster_306 Pantoea
Cluster_254 Myxococcales
Cluster_71 Clostridium sensu stricto 1
Cluster_36 Bacteroides
Cluster_22 Staphylococcus
Cluster_65 Pseudomonas
Cluster_8 Bifidobacterium
Cluster_19 Bradyrhizobium
Cluster_110 Spirosoma
Cluster_101 Polaromonas
Cluster_28 Xanthobacteraceae
Cluster_167 Acinetobacter
Cluster_140 Bacillus
Cluster_180 [Agitococcus] lubricus group
Cluster_234 Gammaproteobacteria genus
Cluster_154 Leucobacter
Cluster_134 Polaromonas
Cluster_35 Bacillus
Cluster_63 Rhodocyclaceae C39
Cluster_372 Lachnospiraceae NK4A136 group
Cluster_61 Lachnospiraceae
Cluster_245 Bacteroides
Cluster_187 Multi-affiliation
Cluster_432 Bacillus
Cluster_107 Thermoactinomyces
Cluster_47 Burkholderiaceae
Cluster_118 Rhodoferax
Cluster_21 Intrasporangiaceae
Cluster_337 Unknown
Cluster_43 Streptococcus
Cluster_85 Actinoplanes
Cluster_73 Solirubrobacter
Cluster_266 Blautia
Cluster_44 Ruminococcus 2
Cluster_241 Diplorickettsiaceae

**Fig. S5**. Abundance of 16S rRNA gene copies of taxonomic units detected in second set of blood samples (n=42) that significantly correlated with the serum levels of zonulin. ρ, Spearman's rank correlation coefficient; *P*, P value of the Kendall's rank correlation. Taxa that resulted significantly correlated with zonulin also from the analysis of the first set of blood samples are indicated in bold and red color.

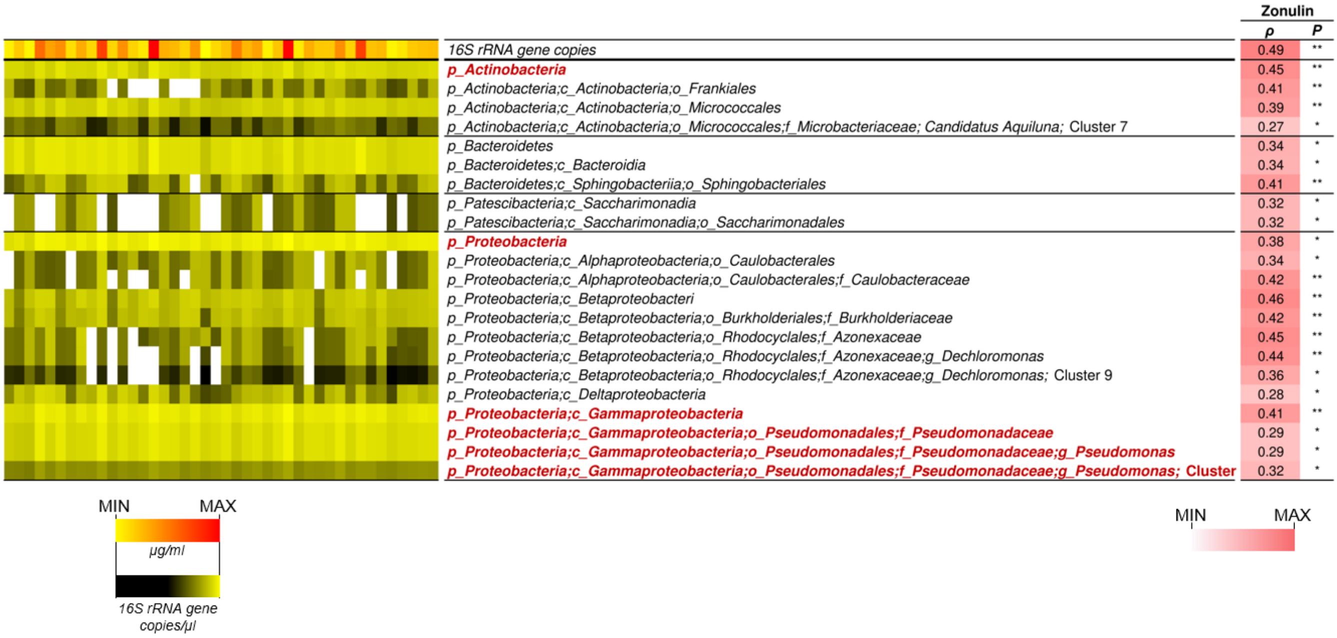
